# Global cis-regulatory landscape of double-stranded DNA viruses

**DOI:** 10.1101/2025.07.20.665756

**Authors:** Tommy Henry Taslim, Joseph Alexander Finkelberg, Susan Kales, Luis Soto-Ugaldi, Benedetta D’Elia, Berkay Engin, George Muñoz-Esquivel, Elvis Morara, Jacob Purinton, Harshpreet Chandok, Jaice Theodore Rottenberg, Rodrigo Castro, Lucia Martinez-Cuesta, Matias Alejandro Paz, Ryan Tewhey, Juan Ignacio Fuxman Bass

**Affiliations:** Molecular Biology, Cell Biology and Biochemistry Program, Boston University, Boston, MA 02215, USA; Bioinformatics Program, Boston University, Boston, MA 02215, USA; The Jackson Laboratory, Bar Harbor, ME 04609, USA; Tri-institutional Program, Computational Biology and Medicine, New York, NY 10065, USA; Biology Department, Boston University, Boston, MA 02215, USA; Escuela Profesional de Genetica y Biotecnologia, Facultad de Ciencias Biológicas, Universidad Nacional Mayor de San Marcos, Lima 15081, Peru

## Abstract

Most double-stranded DNA (dsDNA) viruses use the host transcriptional machinery to express viral genes for replication and immune evasion. This is mediated by viral cis-regulatory elements (CREs) regulated by host and viral transcription factors (TFs). Although some viral CREs and their regulatory mechanisms have been determined, most remain unidentified. Here, we used massively parallel reporter assays to identify ∼2,000 CREs across 27 dsDNA viruses from the Adenovirus, Herpesvirus, Polyomavirus and Papillomavirus families. Viral genomes have a higher CRE density than the human genome, with most viral CREs having promoter-like features and overlapping protein coding sequences. Using saturation mutagenesis and machine learning models, we report viral CRE regulators, including SP, ETS, bZIPs, and TFs acting downstream of signal-activated pathways. Altogether, we present a comprehensive functional CRE map of human-infecting dsDNA viruses that serves as a blueprint for further studies in viral regulation, reactivation, evolution, and viral vector design.

## Introduction

Double-stranded DNA (dsDNA) viruses cause various diseases, including acute infections and those associated with viral persistence, such as autoimmunity, dementia, and cancer [1–4]. These viruses have complex replication cycles and often remain latent until cellular or environmental signals promote their reactivation [5]. Viral cis-regulatory elements (CREs) are key to these processes by regulating the expression of viral genes necessary to replicate, remain latent, or evade immune responses [6]. These CREs are controlled by the host transcriptional machinery and, therefore, rely on host, in addition to viral, transcription factors (TFs). Host TFs, acting downstream of various signaling pathways, enable viral CREs to sense and respond to immune activation, stress, and cell proliferation and differentiation, ensuring that viral gene expression adapts to specific cellular conditions [6]. Immune responses, for example, can reactivate viruses like polyomaviruses and adenoviruses by activating pathways that impinge on NF-kB and AP-1 which activate viral promoters [6,7]. Similarly, stress signals such as hypoxia and glucocorticoids can trigger reactivation of latent herpesviruses (e.g., HHV-1 and HHV-8) by activating CREs through HIF1A, EPAS1, and the glucocorticoid receptor [8–10].

Identifying viral CREs and determining their regulatory mechanisms is crucial for understanding replication strategies, reactivation triggers, and to develop targeted therapeutic approaches against viral infections, similar to shock-and-kill to cure HIV [11]. Despite nearly four decades of viral CRE studies, only a small fraction of the viral CRE landscape has been explored. For instance, TF binding has only been determined for ∼140 CREs of dsDNA viruses, involving <500 TF-CRE interactions (**Table S1**). Transcription initiation studies, however, have identified 190 and 322 transcription start sites (TSSs) in HHV-1 and HHV-4, respectively [12,13]. This represents an average of 1-2 TSSs per kb, much higher than the 0.15 CREs per kb characterized for TF binding or regulation, even in these highly studied viruses (**Table S1**). CRE characterization from less studied viruses is even more sparse.

Here, we used massively parallel reporter assays (MPRAs) tiling through the genomes of 27 viruses from the herpesvirus, adenovirus, papillomavirus, and polyomavirus families and identified ∼2,000 viral CREs, greatly expanding the repertoire of characterized viral CREs by ∼14-fold. Many of these CREs are promoter-like elements near TSSs and are regulated by host TFs, including SP, ETS factors, bZIPs, and TFs acting downstream of signal- or ligand-activated pathways. However, most viral CREs could not be classified into ENCODE CRE classes and overlap with coding sequences, indicating that many viral CREs are different from human ones. Using this viral CRE resource, we describe regulatory differences between viral families, types, and isolates and the drivers of these differences. Finally, we leveraged our TF predictions to engineer an adenoviral vector with reduced background activity, increasing transgene inducibility. Altogether, we present a comprehensive functional CRE map of human-infecting viruses that can guide future studies on viral regulation, reactivation, evolution, and vector design.

## Results

### Identification of CREs from dsDNA viruses

To identify CREs of human-infecting dsDNA viruses, we performed MPRAs tiling through the genomes of 8 herpesviruses (HHV-1 to HHV-8), 8 adenoviruses (types 1, 3, 4, 5, 7, 11, 14, 37), 8 human papillomaviruses (HPVs 1, 2, 5, 6b, 11, 16, 18, and 52), and 3 polyomaviruses (BKPyV, JCPyV, and MCPyV) (**Table S2**). Each genome was tested using 200 bp tiles with 150 bp overlapping windows, and probed in both orientations (positive and negative) (**Figure 1A**). This library contained 66,026 viral tiles, as well as positive (91 sequences active in MPRAs across cell lines) and negative (250 shuffle sequences, 1,876 sequences overlapping human ORFs, and 506 sequences without MPRA activity) controls (**Table S3**). Each tile was associated with an average of 324 barcodes to increase the sensitivity and confidence in activity measurements. MPRAs were conducted in 3-5 replicates in six cell lines representing different model systems for the viruses tested: GM12878 (lymphoblastoid), Jurkat (T cell), MRC-5 (fibroblast), A549 (epithelial), HEK293T (epithelial-like), and K562 (hematopoietic) cells. These experiments passed stringent quality control metrics, including number of barcodes per tile, number of read counts, correlation between replicates, and activity levels of control sequences (see **Figure S1A-C** and **STAR Methods**). Control sequences were used to establish thresholds corresponding to a 1% false discovery rate in each cell line for identifying active viral tiles (**Figure S1C**). This resulted in 5,275-6,896 active tiles per cell line, which displayed a broad range of activities spanning 3 orders of magnitude (**Figure S1D**).

**Figure 1:**
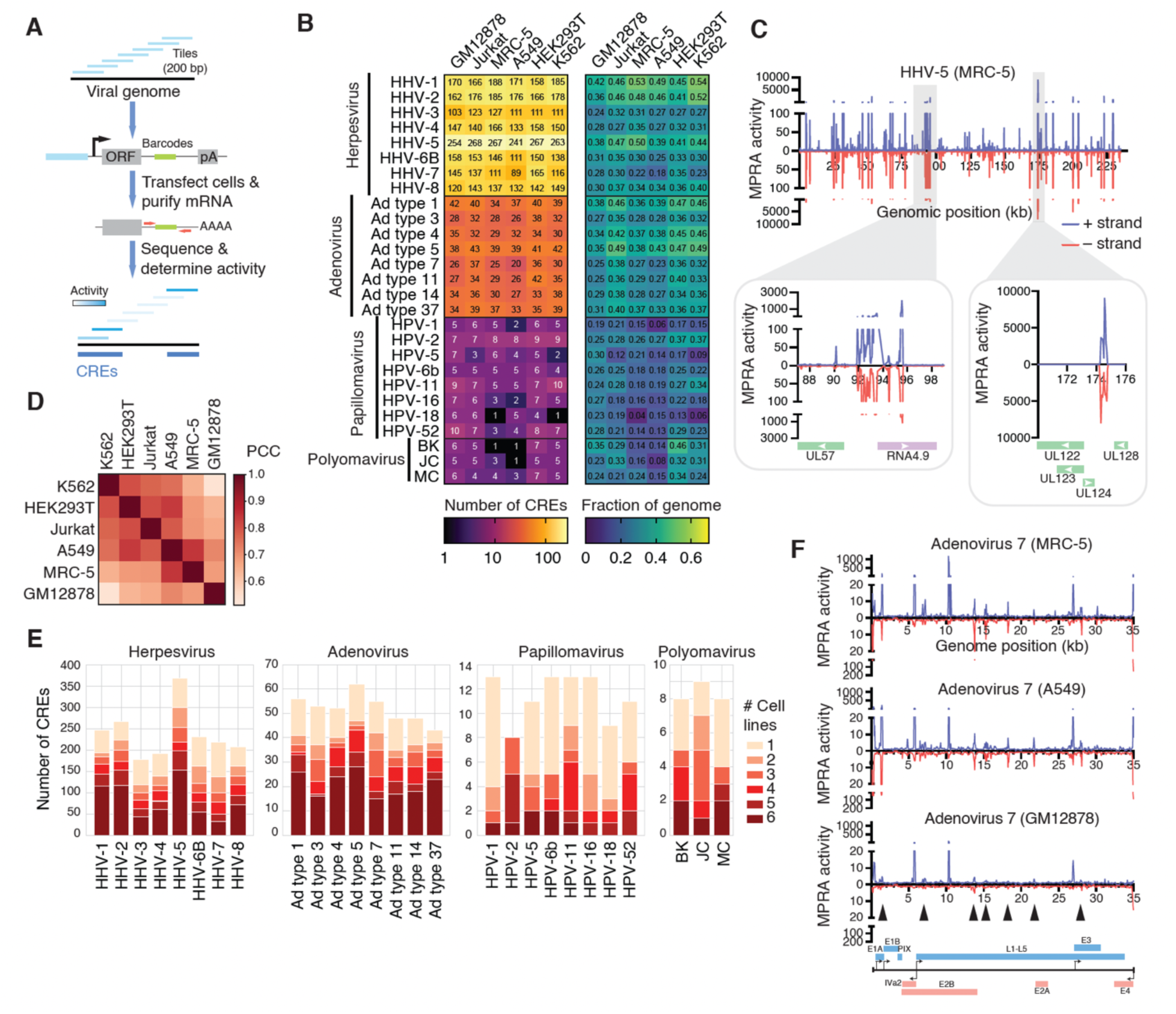
Identification of CREs across the genomes of dsDNA viruses. (A) Schematic of MPRAs tiling through the genomes of 27 different dsDNA viruses. Tiles consistent on 200 bp sequences with offsets of 50 bp. Sequences were tested in both orientations. (B) Number and fraction of the genome covered by active CREs in each cell line for each virus tested by MPRAs. (C) MPRA activity map for HHV-5 in MRC-5 cells. In blue and red is the activity when tiles were in positive and negative strand orientation, respectively. Insets show examples of two highly active regions. (D) Correlation (Pearson correlation coefficient) in activity between cell lines across tiles active in at least one cell line. (E) Number of CREs active in 1 to 6 cell lines for each virus tested. (F) Activity maps for adenovirus 7 in MRC-5, A549, and GM12878 cells. Arrows indicate CREs for which activity was lost in GM12878 cells.

To define genomic regions with CRE activity, we merged overlapping active tiles (**Figure S1E**). This resulted in 1,444-1,687 active CREs per cell line, and 2,061 CREs considering all cell lines and viruses (**Table S4**). We successfully recovered 94% of CREs reported in the literature (**Table S1**), highlighting the high coverage of our MPRA experiments. As expected, the number of CREs correlated with genome size, with herpesviruses having the largest number of CREs and polyomaviruses and papillomavirus having the smallest (**Figure 1B**). On average, this corresponds to 0.97 CREs/kb per cell line, substantially higher than the 0.07 CREs/kb per cell type predicted in the human genome and similar to the density observed in human promoter regions (**Figure S1F**) [14]. Altogether, this is consistent with dsDNA viral genomes containing a much higher density of genes than eukaryotic genomes, which require independent transcriptional control.

Human cytomegalovirus (HHV-5), whose genome is ∼235 kb, has the largest number of CREs (267 CREs in MRC-5 cells) among the viruses tested (**Figure 1C**). Notably, 9 CREs had activity exceeding 1,000-fold relative to negative controls, including a 650 bp region covered by consecutive 10 tiles, all with activities above 1,000 (**Figure 1C**). This region is upstream of immediate early TFs IE1/IE2, essential for HHV-5 replication [15,16]. We also identified a highly active ∼2kb region between UL57, encoding a single-stranded DNA binding protein, and the long noncoding RNA4.9, which regulates viral DNA replication [17]. This shows that, even with small genomes, viruses can harbor large CRE regions.

### Viral CRE activity is mostly conserved across cell lines

Different cell types express distinct TF sets and, therefore, viral CRE activity may differ across cell lines. Despite potential differences in TF expression, we observed moderate to high activity correlation (0.48-0.83) between cell lines, with GM12878 and MRC-5 being the most different to other cell lines (**Figures 1D** and **S2A-B**). Most CREs are active across all cell lines (**Figures 1E-F**), suggesting they are generally regulated by core constitutive TFs rather than cell type-specific ones. This is particularly evident for herpesvirus and adenovirus CREs, where only 25% of CREs are cell line-specific. Universally active CREs have higher activity and are larger than CREs active in a subset of cell lines (**Figure S2C**). However, we also observed multiple instances where CRE activity was variable across cell lines (**Figures 1F** and **S2D**). For polyomavirus and papillomavirus, specificity may reflect differences in regulation across cell types, especially for CREs other than the main transcriptional control elements (non-coding control region - NCCR for polyomavirus, and long control region - LCR for papillomavirus). Interestingly, CRE activity specificity is not directly correlated with viral tropism. This is consistent with tropism being defined mainly by cellular receptors for entry and the cellular machinery required for replication and virion assembly [5].

### Classification of viral CREs

Given the compactness of viral genomes, we hypothesized that many CREs would overlap with or be near TSSs, potentially acting as gene promoters. Using TSS positions from 5’ end sequencing of viral transcriptomes determined for HHV-1 [12], HHV-3 [18], HHV-4 [19], and HHV-8 [20] we found that tiles containing TSSs exhibited significantly higher MPRA activity than those lacking TSSs (**Figure 2A**). We also observed that, across most viruses, viral CREs are closer to annotated TSSs (<250 bp away from the 5’ end of annotated gene coordinates) than expected by chance when randomizing CRE positions in the genome (**Figures 2B**). Similarly, we found that most TSSs have at least one CRE within 250 bp (**Figure 2C**). These findings suggest that many CREs may function as promoter elements.

**Figure 2:**
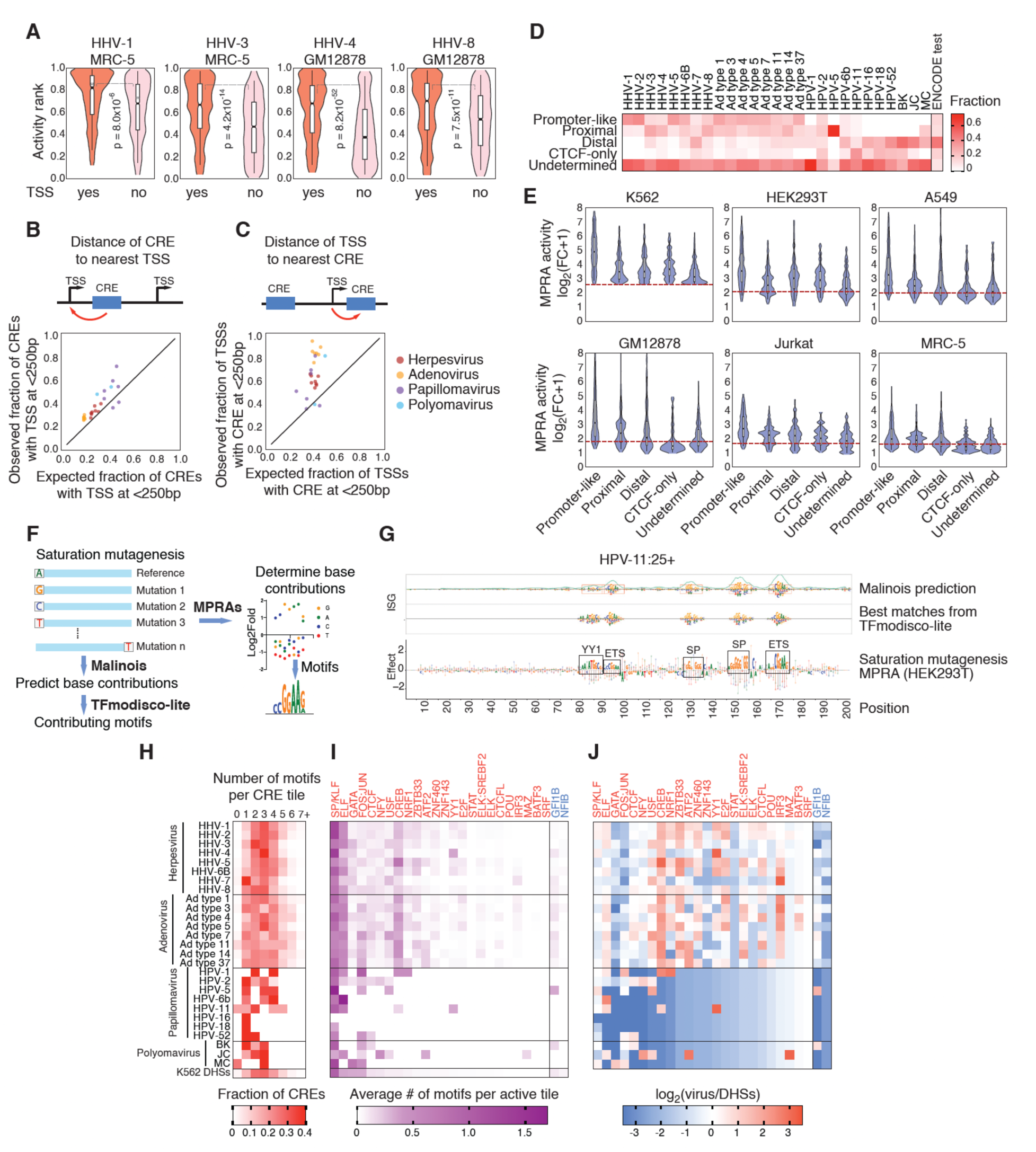
Classification of viral CREs. (A) Distribution of MPRA activity ranks for HHV-1, HHV-3, HHV-4, and HHV-8 tiles that contain experimentally determined TSSs. Violin plots are shown together with box plots (median = solid black line, quartiles = box). Statistical significance determined by Kolmogorov-Smirnov test. (B) Observed versus expected distance of CREs active in HEK293T cells to the closest annotated transcription start site (TSS). Scatter plot of the observed versus expected fraction of CREs whose midpoint is located within 250 bp of an annotated TSS for each virus tested. (C) Observed versus expected distance of annotated TSSs to the closest active CRE in HEK293T cells. Scatter plot of the observed versus expected fraction of annotated TSSs located within 250 bp of the midpoint of an active CRE for each virus tested. (D) Fraction of active tiles per virus classified by CRE-class as promoter-like, proximal, distal, CTCF-only, or undetermined. Human DHSs active in K562, HepG2, or SK-N-SH cells are shown for comparison. (E) MPRA activity of tiles from each tile class active in K562 cells. For each tile, the maximum activity between the positive and negative strand is considered. The dashed red lines indicate the threshold to consider a tile active in the corresponding cell line. Violin plots are shown together with box plots (median = solid black line, quartiles = box) for each class of element in each cell line. (F) Schematic of saturation mutagenesis experiments and analyses. Each base pair of an active tile is substituted for the alternative 3 bases and activity is determined using MPRAs. Base contributions to activity are used to derive activity contributing motifs. Similarly, base contribution can be predicted using Malinois and TFmodisco-lite can be used to determine motifs that contribute to activity. (G) Example of an HPV-11:25+ tile showing the base contributions predicted using Malinois trained in K562 cells, and the best matches derived from TFmodisco-lite. Below are base contributions from the MPRA saturation mutagenesis experiments in HEK293T cells. (H) Distribution of number of motifs per active CRE tile across different viruses and human DHSs predicted to be active in K562 cells. (I) Average number of motifs per active tile derived from TFmodisco-lite and collapsed into single TF motifs. Activating and repressing motifs are shown in red and blue, respectively. (J) Average number of motifs per active tile relative to their frequency in human DHSs active in K562 cells.

We next sought to identify active viral tiles with sequence features resembling different types of human CREs defined by the ENCODE project (promoter-like, proximal, distal, CTCF-only) [21]. Previous studies have shown that promoter and distal sequences have distinct TF motif compositions, with some motifs imparting promoter or enhancer features to human CREs [22,23]. We hypothesized that the core 200 bp sequence would be sufficient to classify human and viral CREs in one of four ENCODE classes based on TF motif presence. To test this, we used the pre-trained Malinois convolutional neural network that predicts MPRA activity and has learned TF motif syntax in three human cell lines: K562, HepG2 and SK-N-SH [14]. We developed CRE-Class, a model based on Malinois’ convolutional layers that predicts CRE classes directly from DNA sequence. Promoter-like and distal human CREs exhibited distinct dinucleotide compositions (**Figure S3A**). To enforce CRE-Class to learn characteristic TF motifs and prevent just learning dinucleotide composition, we added trinucleotide-shuffled artificial CRE sets (see **STAR Methods**). Assessment of CRE-Class on the test set of human CREs confirmed that distal and promoter-like sequences could be distinguished with high recall and precision. However, the model struggled to distinguish CTCF-only from distal sequences, and proximal elements were often classified as promoter-like or distal CREs (**Figures S3B-C**). This aligns with the ENCODE CRE classification, where CTCF-only and distal classes are interchangeable in certain cell types, and proximal elements are classified according to a heuristic distance to TSS threshold [21]. CRE-Class successfully distinguished real CRE classes from the shuffled ones. Examination of contribution scores confirmed that the model learned actual motifs, not just base composition differences between promoter and distal CREs (**Figure S3D**). Thus, CRE-Class can accurately classify human CREs into promoter-like and distal classes based on TF motif composition.

Driven by the high precision and recall of CRE-Class and Malinois’ ability to predict K562 MPRA activity of viral sequences (**Figures S3E-F**), we applied CRE-Class to viral CREs. Nearly half of active viral tiles matched shuffled CRE classes (mostly Promoter-like shuffled) and were deemed “undetermined”, compared to only 9.7% of the human CRE test set (**Figure 2D** and **Table S5**). We observed a significant enrichment of promoter-like or proximal CREs and a depletion of distal CREs in most herpesviruses (except HHV-7) and adenoviruses relative to human DHSs (**Figure 2D**). This pattern was absent in papillomavirus and polyomavirus CREs whose main transcriptional control elements are known to be regulated by TFs typically enriched in distal CREs, such as bZIP factors (**Table S1**). Consistent with previous studies on human CREs [22], promoter-like CREs are more active than distal, proximal, and CTCF-only CREs across viruses and were more broadly active across cell lines (**Figures 2E** and **S3G**). Altogether, this shows an enrichment of highly active promoter-like elements, consistent with the compactness of viral genomes.

### Multiple human TFs contribute to viral CRE activity

To identify TFs regulating viral CRE activity, we performed *in silico* saturation mutagenesis using Malinois trained on K562 cells. We predicted the activity effect of all possible single base substitutions in each active viral tile and human DHSs in K562 cells. Using TFmodisco-lite, we identified motifs contributing to activity across viral tiles and human DHSs, which were then matched to known motifs using TomTom [24] (**Figure 2F** and **Table S6**). To validate these predictions, we conducted MPRA saturation mutagenesis on 39 active tiles mutating each base to the three alternative bases, and observed strong concordance with model predictions (**Figure 2G**). In total, TFmodisco-lite identified 45 motifs (43 activating and 2 repressing) across all human and viral active tiles, including single and composite motifs (**Figure S4A**). These motifs were collapsed into 26 single TF motifs.

While active human DHSs contained a median of 2.88 motifs per 200 bp, viral tiles showed median values ranging from 1.1 and 2.9, depending on the virus (**Figure 2H**). The higher number of motifs in human DHSs may reflect the precise centering of DHS peaks over regions of maximal chromatin accessibility. In contrast, viral tiles are positioned systematically across the genome, leading to tiles that partially overlap with CREs having fewer motifs than those at the center. Herpesviruses and adenoviruses often have more activity-contributing motifs than papillomaviruses and polyomaviruses, although there is variability within viral families. The most prevalent motifs across viruses correspond to activating SP/KLF, ELF, and CREB motifs (**Figure 2I**). This is consistent with studies that identified multiple SP and bZIP binding sites across viral CREs (**Table S1**). Interestingly, ETS factors account for only 2% of documented binding/regulatory events, whereas Malinois predicts they account for 15% of active binding sites. Given that ETS factors are influenced by pathways like MAPK/ERK, PI3K/Akt, and JAK/STAT, which promote the reactivation of certain latent viruses [25], this suggests that ETS factors play a larger role in viral regulation and response to cellular states than previously reported.

Several motifs were enriched in viral CREs compared to human DHSs. For example, activating CREB, YY1, IRF3, NRF1, ZBTB33, and E2F were enriched across herpesviruses and adenoviruses relative to human DHSs (**Figure 2J**). These TFs act downstream of diverse signaling pathways, including MAPK/ERK, PI3K/Akt, cGAS-STING, wnt/b-catenin, and PKA pathways, that signal different cellular states known to trigger viral reactivation or immune evasion responses [6,26,27]. Conversely, activating FOS:JUN and GATA motifs, and repressive GFI1B and NFIB motifs, were depleted in CREs from most viruses relative to human DHSs. Some of these differences can be attributed to differing proportions of CRE classes between human and viral CREs. Nevertheless, we also found differences between human and viral CREs from each class (**Figure S4B**). For example, viral promoter-like CREs are enriched in CREB, IRF3, and USF motifs relative to human promoter-like CREs. Further, there is significant variability in motif representation in each type of CREs across viruses, which, together with a different proportion of CREs types, suggests different regulatory strategies between viruses even within the same viral family.

### Most viral CREs overlap with protein coding sequences and are classified as undetermined

Given the enrichment of promoter-like elements, we anticipated that many viral CREs would be located in intergenic regions. Indeed, we found an overall enrichment of CREs in intergenic regions and a depletion in coding regions relative to randomized CRE positions across the viral genomes (**Figure 3A**). However, because viral genomes contain a high density of protein coding regions, 35-93% of the CRE tiles overlap with coding regions, depending on the virus. This represents a stark contrast to human CREs, of which 2.4% overlap with coding sequences as predicted by Malinois. Interestingly, TF motifs predicted by Malinois to contribute to activity cover 10.5% of viral codons and are generally enriched in intrinsically disordered regions (**Figure 3B**). These regions are generally less evolutionarily constrained than structured domains, which could reduce constraints for simultaneous evolution of CREs and protein coding sequences.

**Figure 3:**
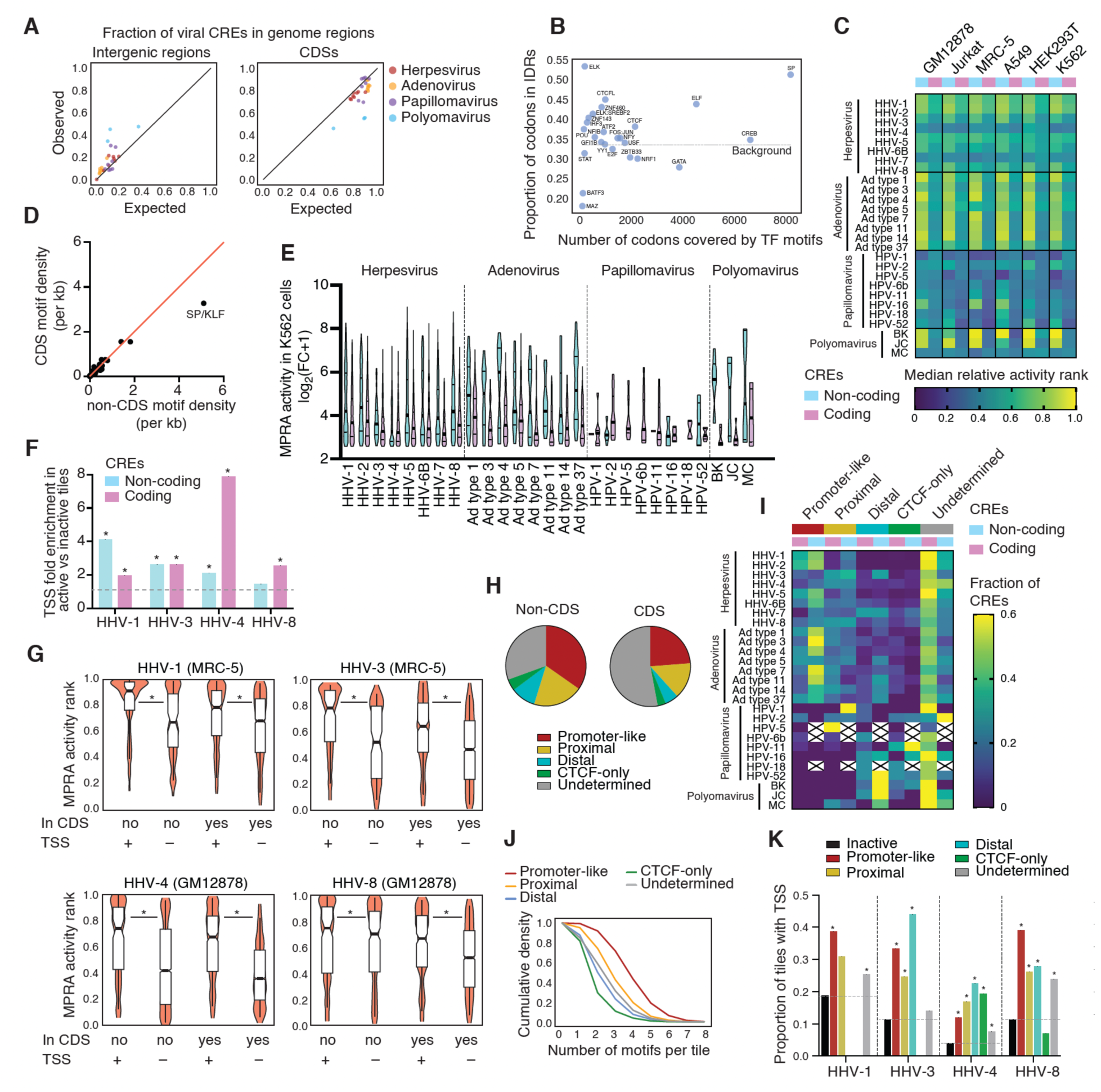
Most viral CREs overlap coding sequences. (A) Scatter plot of the observed versus expected fraction of CREs whose midpoint is located in intergenic (left) or coding (right) regions for each virus tested. (B) Proportion of codons in intrinsically disordered regions (IDRs) intersecting with motifs predicted using Malinois versus number of codons intersected by each TF across all viruses tested. (C) Heatmap of median MPRA activity rank of active tiles overlapping coding and non-coding regions per virus and per cell line. Crosses indicate data not available. (D) Scatter plot of the density of TFs motifs for CRE tiles in coding versus non-coding regions. (E) MPRA activity distribution in K562 cells for tiles overlapping coding and non-coding regions for each virus. Violin plots are shown with the median (solid line) and quartiles (dotted lines). (F) Enrichment of tiles containing experimentally determined TSSs for active relative to non active tiles in non-coding and coding regions. (G) Distribution of MPRA activity ranks for HHV-1, HHV-3, HHV-4, and HHV-8 tiles that contain or not CAGE peaks or annotated 5’ transcript ends. Violin plots are shown together with box plots (median = solid black line, quartiles = box). *p<0.05 by Kolmogorov-Smirnov test. (H) Fraction of active tiles of different classes among the CREs overlapping with coding and non-coding regions across all viruses. (I) Fraction of active tiles in coding and non-coding regions corresponding to different CRE classes for each virus. Crosses indicate data not available. (J) Cumulative density of the number of motifs per active tile for each CRE class. (K) Proportion of tiles containing experimentally determined TSSs. *<0.05 by one-tailed proportion comparison test.

CRE tiles in coding regions are generally less active than those in non-coding regions, especially for adenovirus and polyomavirus, and to a lesser extent, herpesvirus (**Figure 3C**). This is likely due to a lower density of activating SP/KLF motifs in CREs in coding regions (**Figure 3D**). However, individual tiles displayed a wide distribution of activity, with many CRE tiles in coding regions being highly active (**Figure 3E**). Additionally, we observed a similar enrichment for TSSs for CRE tiles in coding and non-coding regions (**Figure 3F**). Finally, tiles in coding regions containing TSSs were more active than those without TSSs, similar to CRE tiles in non-coding regions (**Figure 3G**). Altogether, this shows that CREs in coding sequences are functional and may be important drivers of viral gene expression.

Coding regions are enriched in undetermined tiles, and depleted of promoter-like, proximal, and distal tiles (**Figure 3H-I**). These undetermined tiles have similar dinucleotide frequencies and relative proportion of contributing TF motifs to promoter-like viral tiles, but fewer motifs per tile and overall lower activity (**Figures 2E**, **3J**, **S3H** and **S4B**). However, 20% of undetermined tiles had activity levels above 10-fold relative to negative controls and were enriched for TSSs, similar to other types of viral CREs (**Figure 3K**). This, along with the large fraction of undetermined tiles, suggests that undetermined tiles may be major contributors to viral transcriptional regulation.

### Identification and characterization of polyomavirus CREs

We determined the MPRA activity across the genomes of three polyomaviruses BKPyV (MM), JCPyV (MAD-4), and MCPyV (MKL-1) in HEK293T cells, which expresses the SV40 large T-antigen that controls viral gene expression and replication (**Figure 4A**). In all three viruses, the most active region corresponded to the NCCR, a bidirectional promoter controlling early and late gene expression and that drives type-specific pathogenesis [28–30]. Previous studies have identified multiple TFs binding or regulating NCCR activity of BKPyV and JCPyV, while little is known about MCPyV NCCR regulation (**Table S1**) [6,31,32]. These studies were based on perturbations of known motifs or were restricted to TFs for which antibodies were available. To determine the TFs that contribute to the NCCR activity in an unbiased manner, we performed saturation mutagenesis MPRAs of NCCR tiles, identifying multiple activating and a few repressive motifs (**Figures 4B-C**). This included previously reported TF interactions (AP-1, NF-kB, NFI, AP-2, SP1, and large T-antigen) and novel TF motifs (**Table S1**). Most motifs, except for activating AP-1 sites and one repressive NFI site, were not shared across the three viruses, indicating distinct regulatory NCCR architectures, consistent with the low sequence conservation in these regions [33]. For example, SP/KLF only activates BKPyV and MCPyV, whereas NFI only activates BKPyV and JCPyV (**Figures 4B-C** and **S5A-B**). Interestingly, MCPyV has been split into two classes based on a 25 bp repeat in the NCCR, with strains containing two repeats having a potentially more active NCCR than those with one repeat [34]. We show that this region contains an active SP/KLF site (**Figure S5B**). These SP sites seem to be non-redundant, and may function additively or cooperatively in homotypic clusters, as single nucleotide perturbations of any of the 3 SP/KLF sites in tile 102+ reduced activity.

**Figure 4:**
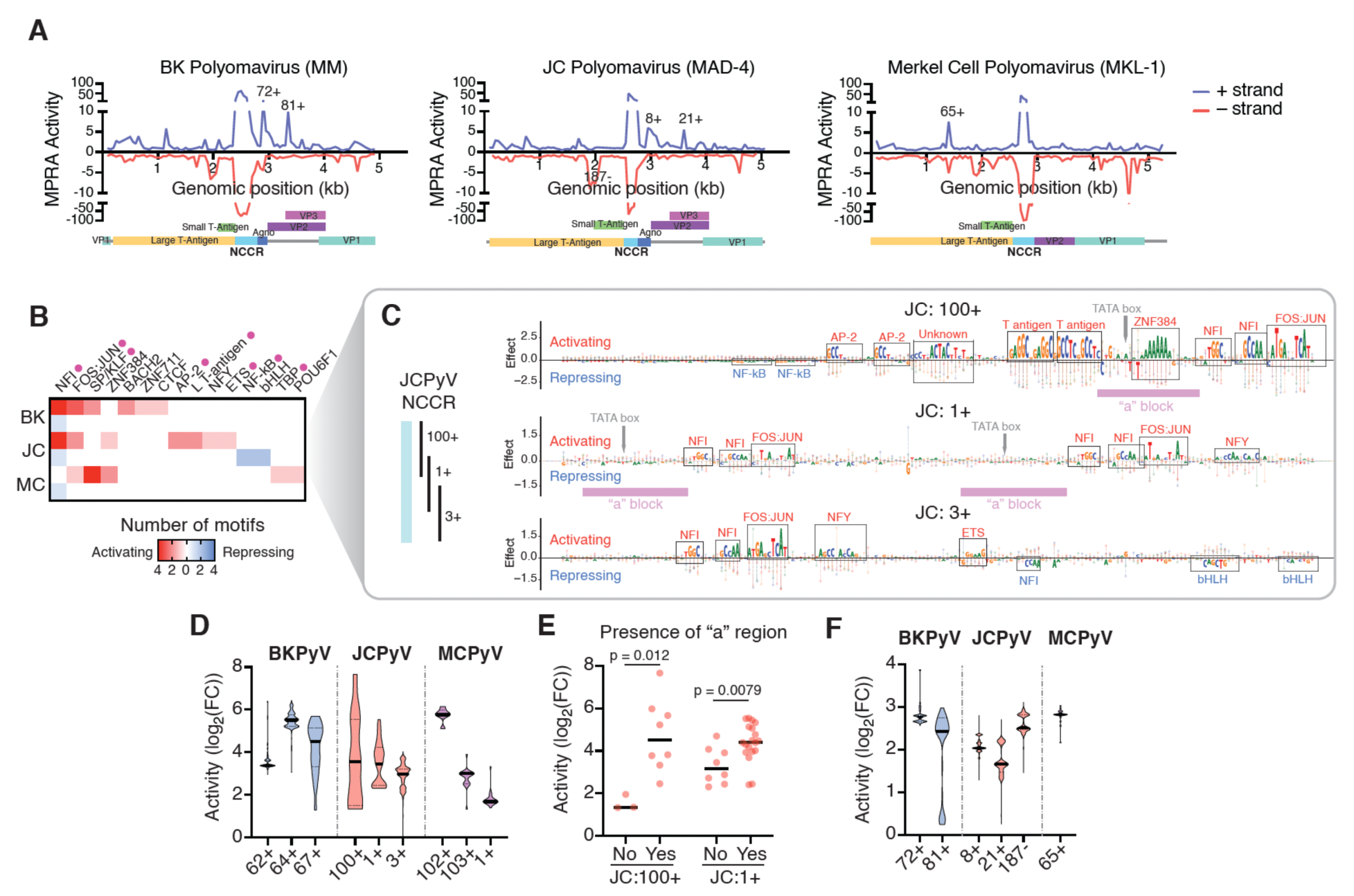
Characterization of Polyomavirus CREs. (A) MPRA activity maps for BK, JC, and Merkel Cell Polyomaviruses in HEK293T cells. Genome maps depicting the main genes and the noncoding control region (NCCR) are shown below. (B) Number of activating or repressing motifs for each TF in the NCCRs of the BK, JC, and Merkel Cell polyomaviruses obtained from MPRA saturation mutagenesis experiments in HEK293T cells. (C) Example of MPRA saturation mutagenesis in HEK293T cells for tiles 100+, 1+ and 3+ that cover the JC polyomavirus NCCR. Activating and repressive motifs are indicated in red and blue, respectively. (D) Violin plots of the activity measured by MPRAs in HEK293T cells for different isolates from BK, JC, and Merkel Cell polyomaviruses across tiles that cover the NCCR. The median and quartiles are indicated by bold solid and thin dashed lines, respectively. (E) Activity comparison between JC polyomavirus isolates that include the “a” region and those that do not, for tiles 100+ and 1+. Statistical significance determined by Mann-Whitney’s U test. (F) Violin plots of the activity measured by MPRAs in HEK293T cells for different isolates from BK, JC, and Merkel Cell polyomaviruses across different active tiles. The median and quartiles are indicated by bold solid and thin dashed lines, respectively.

Among the novel motifs, we found an activating poly-A motif -matching ZNF384-downstream of the TATA box in JCPyV and MCPyV. While this sequence is also present in the BKPyV NCCR, it does not contribute to MPRA activity (**Figure S5A**), possibly due to a positional effect or to functional redundancy with other TFs. We also identified a novel activating ZNF711 motif specific to BKPyV, which we confirmed using enhanced yeast one-hybrid assays (**Figure S5C**), an activating NFY motif in JCPyV, two repressive bHLH sites in JCPyV, and several unidentified activating and repressive motifs.

The NCCR sequence is highly variable, composed of different sequence blocks with insertions, deletions, nucleotide variants, and differences in copy number, often linked to strain-specific pathogenesis [33,35]. For example, in JCPyV, these blocks (sized 18-69 bp) are called a-f with their combination and repeats defining NCCR types. To assess activity variability and contribution of different blocks, we performed MPRAs in HEK293T cells using sequences from multiple polyomavirus isolates (342 from BKPyV, 510 from JCPyV, and 22 from MCPyV) (**Table S7**). MCPyV tiles showed limited activity variability (**Figure 4D**), consistent with MCPyV being a more recent and less variable virus than JCPyV and BKPyV [34]. Despite tiles 62+ and 64+ in BKPyV and 3+ in JCPyV being highly variable in sequence, they display a marked conservation in activity, with the interquartile range being less than 2-fold (**Figure 4D**). Other regions instead, such as tiles 67+ from BKPyV, and tiles 100+ and 1+ from JCPyV have a much higher activity variability, with interquartile ranges between 3.5 and 16.6 fold. The major driver of activity variability of tiles 100+ and 1+ from JCPyV is the presence of an “a” block which contains the TATA box and the ZNF384 motif (**Figure 4E**). Interestingly, tile 1+ from JCPyV MAD-4 strain, which has 2 copies of block “a”, showed no bases within “a” contributing to MPRA activity. This can be due to redundancy between both “a” blocks, robustness of motifs difficult to perturb by single nucleotide variants, or a positional effect. In the case of BKPyV:67+, activity variability is driven by differences in multiple TF binding sites, particularly NFI.

In addition to the NCCR, we have also identified regions with transcriptional activity overlapping viral protein coding sequences. For example, we detected a homologous CRE in the Agno coding region in BKPyV (tile 72+) and JCPyV (tile 8+) (**Figure 4A**), whose activity is highly conserved across BKPyV and JCPyV isolates (**Figure 4F**), consistent with the overall sequence conservation. We also identified non-homologous CREs in VP2 of BKPyV (tile 81+) and JCPyV (tile 21+) with moderate activity conservation across isolates, and CREs with moderate/high activity conservation within the large T-antigen of JCPyV (tile 187-) and MCPyV (tile 65+) (**Figures 4F and S5D-E**). These CREs were classified as distal or CTCF-only by CRE-Class, consistent with a previous study showing these regions in SV40 Polyomavirus have the enhancer H3K4me1 mark [36]. Whether these CREs affect the expression of early or late genes, remains to be determined.

### Identification and characterization of human papillomavirus CREs

HPVs generally infect epithelial cells and are classified into over 200 types, ranging from low-risk types causing benign warts to high-risk types associated with cancer [5]. To determine how these types are regulated, we determined MPRA activity across the genomes of five low-risk (1, 2, 5, 6b and 11) and three high-risk (16, 18, and 52) types in HEK293T cells. The main HPV CRE corresponds to the LCR, which controls the expression of most viral genes (e.g., E6, E7, L1, etc.); however, activity in this region was only detected in three types (2, 16, and 52), with the highest activity in the high-risk HPV-16 and HPV-52 (**Figure 5A**). This is consistent with previous studies linking high expression levels of oncoproteins E6 and E7 with cancer risk [37,38]. One reason we may have not detected LCR activity in some types is that maximal activity may require viral proteins such as E1 and E2, not expressed in HEK293T cells, and cellular activation [39]. To identify motifs contributing to LCR activity, we performed saturation mutagenesis MPRAs in LCR tiles from HPV-16 and HPV-52. Although we found a shared activating FOS:JUN motif, most motifs differed between these two types (**Figures 5B and S5F-G**). The limited conservation of TF usage is expected given the low sequence homology between the LCRs of HPV-16 and HPV-52. Given the high regulatory variability between HPV types, we next evaluated whether activity varied across isolates from the same type. Most tiles from HPV-16 and HPV-52 LCRs showed low variability, except for a few low and high activity isolates (**Figures 5C-D** and **Table S7**). However, tile 136+ within the central segment of the HPV-16 LCR was highly variable. To identify the nucleotide positions responsible for this variability, we aligned the sequences across isolates and compared their MPRA activity. Although multiple positions showed nucleotide variation, activity changes were mostly explained by variants within the 3’ end of tile 136+, which correspond to positions with high activity contribution based on saturation mutagenesis MPRAs (**Figure 5E**). Most isolates with loss of activity had a C->T variant at position 172 that disrupts the SREBF motif. Other low activity isolates had similar disruptions of this motif (C->T at 170, and T->C at 167). We also found two isolates with substitutions of a G at position 185, which does not belong to a known motif but still contributes to activity. Interestingly, two isolates that substitute a C at position 178, which negatively contributes to activity, resulted in increased activity. These findings show that variants at different positions within viral CREs can lead to extensive variability in activity across isolates.

**Figure 5:**
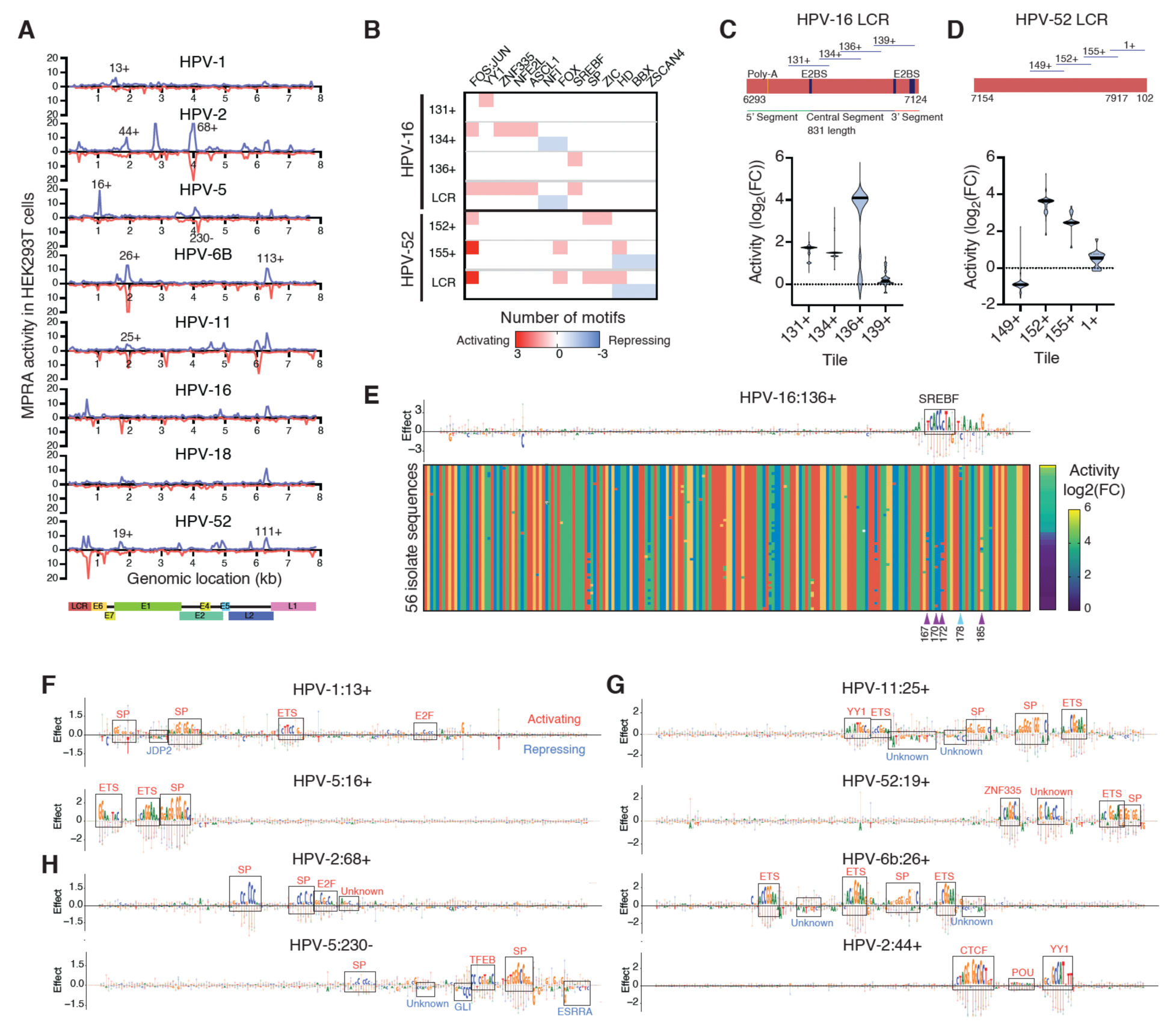
Landscape of CREs across Human Papillomavirus genomes. (A) MPRA activity maps for different HPV strains in HEK293T cells. (B) Number of activating (pink-red) or repressing (blue) motifs for each TF in the tiles overlapping the long control region (LCR) for HPV-16 and -52. Motifs were obtained from MPRA saturation mutagenesis experiments in HEK293T cells. (C-D) Violin plots of the activity measured by MPRAs in HEK293T cells for different isolates from HPV-16 and -52 across tiles that cover the LCR. The median and quartiles are indicated by bold solid and thin dashed lines, respectively. The LCR maps and tiles that cover them are shown above. (E) Activity of 56 HPV-16 isolates for region 136+ of the LCR. The top plot shows the base contributions derived from MPRA saturation mutagenesis in HEK293T cells. The SREBF motif is indicated. The sequence alignment for region 136+ is shown across isolates. A = green, C = blue, G = yellow, T = red. The MPRA activity of each isolate is shown on the right. Purple and teal arrows indicate variants that reduce or increase activity, respectively. (F-H) MPRA saturation mutagenesis experiments in HEK293T cells for regions located in the E7 (F), E1 (G), and E2 (H) genes. TFs contributing to activity are indicated.

In addition to the LCR, we identified multiple CREs across the HPV genomes tested (**Figure 5A**). Some of these CREs have been previously reported. For instance, a CRE upstream of HPV-52 E2 matches a TSS identified by CAGE [40], and a CRE located within HPV-11 E1 was identified by 5’RACE [41]. Further, consistent with 5’RACE or CAGE studies that identified promoters within the E1, E5, and L2 regions of the HPV-16 [42,43], we detected MPRA activity in these regions in several HPV strains. Saturation mutagenesis MPRAs show that many of these CREs contain SP/KLF, ETS, and YY1 motifs often found in promoters, while others were classified as distal or CTCF-only (**Figures 5F-H**). Although some CREs are present in multiple HPV types, the TF binding site identities or locations often differ between types, indicating high regulatory variability beyond the LCR. However, for most non-LCR tiles, we observed high activity conservation between isolates of the same type, consistent with high nucleotide sequence conservation in these coding regions (**Figure S5H**).

### Identification and characterization of Adenovirus CREs

Adenovirus CREs have been hallmark models of viral and eukaryotic transcriptional regulation, with the promoters of E1A, E1B, and late genes being among the first CREs characterized in human cells [44,45]. Adenovirus 5, in particular, has been extensively studied for its impact on cell cycle and its use in viral vector vaccines and cancer therapies [46–48]. Several promoters upstream of E1A, E1B, PIX, E2A/E3, UXP, E4, and L4, and the major late promoter (MLP) have been identified and mechanistically characterized. In MPRAs conducted in HEK293T, A549, MRC-5 and K562 cells, often used as models for adenovirus studies, all these promoters were highly active (**Figure 6A**). Additionally, we identified multiple CREs across the Adenovirus 5 genome, including the L1-L4 late region. A recent study detected an H3K27ac ChIP-seq peak within L2 during early infection, suggesting this may be a promoter or enhancer [49]. Another study found TBP recruitment upstream of L1 and E2B2 [50]. Other CREs in the L1-L4 region had not been detected previously but are highly active, comparable to known Adenovirus 5 promoters, and many were classified as promoter-like or proximal by CRE-Class (**Figure 6A**), suggesting they may correspond to TSSs.

**Figure 6:**
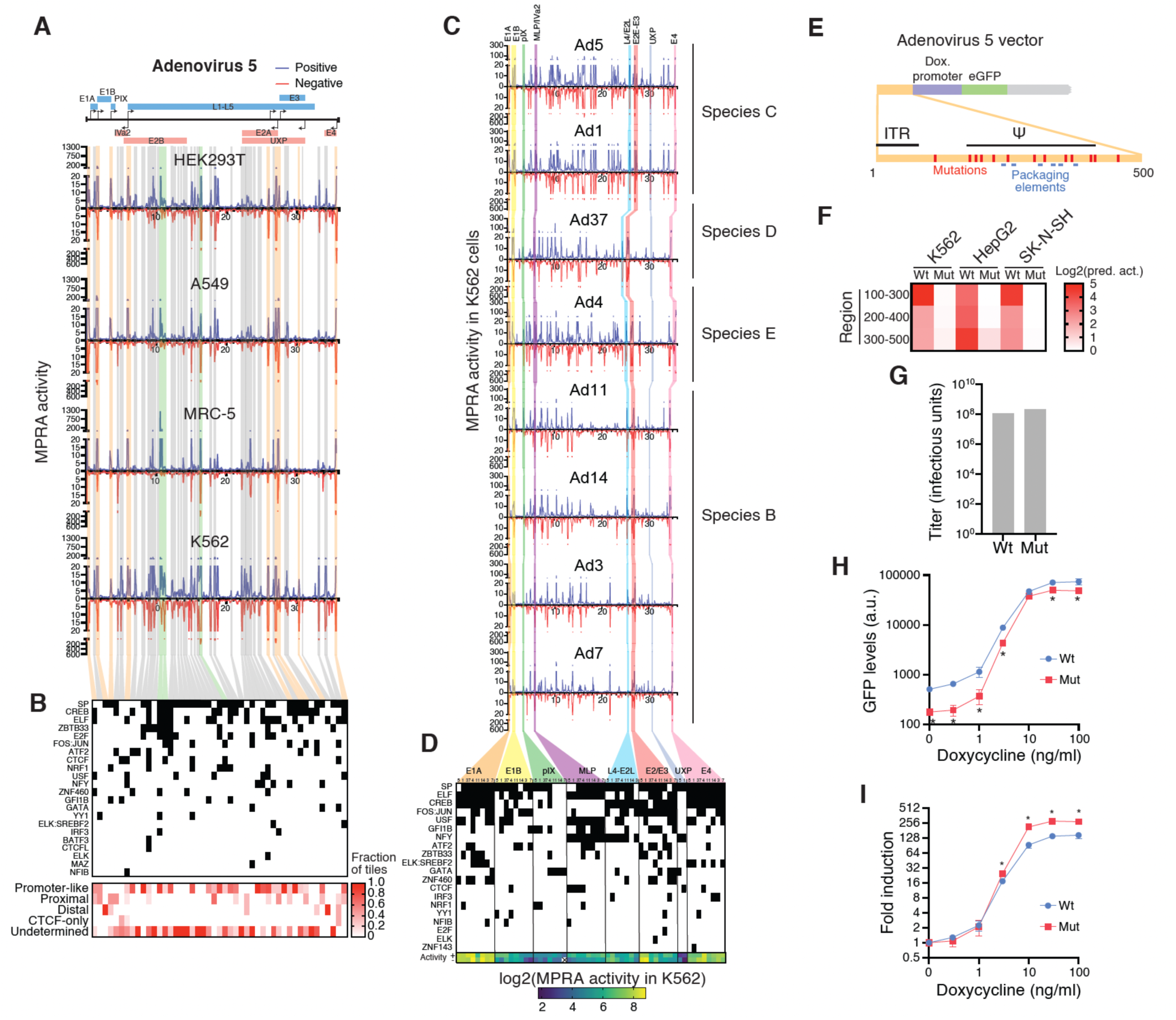
Identification and characterization of adenoviral CREs. (A) MPRA activity map for Adenovirus 5 in different cell lines. In blue and red is the activity when tiles were in positive and negative strand orientation, respectively. Orange shades indicate known promoter regions; green shades indicate other known CREs; gray shades indicate novel CREs. (B) Motifs contributing to MPRA activity in K562 cells for each of the adenovirus 5 CREs. (C) MPRA activity map for different adenovirus strains in K562 cells. In blue and red is the activity when tiles were in positive and negative strand orientation, respectively. Each known promoter is indicated with a different color shade. (D) Motifs contributing to MPRA activity in K562 cells for each of the known promoters across adenovirus strains. (E) Schematic adenovirus 5 vector construct, the mutations introduced to reduce transcriptional activity of the Ψ region, and the regions involved in DNA packaging. (F) Activity of tiles covering the wild type or mutant Ψ sequences of Adenovirus 5 predicted using Malinois trained in K562, HepG2, and SK-N-SH cells. (G) Viral titers obtained with the wild type and mutant Ψ sequence from the Adenobuilder kit. (H, I) Median GFP levels (arbitrary units) determined by flow cytometry (H) and fold-induction relative to vehicle (I) after incubation with different concentrations of doxycycline for 16 hs in cells infected with wild type of mutated Adenovirus 5 vector. *p<0.05 by two-tailed Student’s t-test with Welch correction. Experiments were conducted in biological triplicate.

To determine how Adenovirus 5 CREs are regulated, we used Malinois activity predictions in K562 cells (**Figure 6B**). SP sites were identified in many CREs, including the E1B promoter and the MLP, which have been reported [45,51]. Other predicted interactions, including ATF1/ATF2, CREB1, and JUN with the E2 and E3 promoters, and USF with the MLP, have also been described [44,52,53]. Many known and novel CREs are predicted to be regulated by TFs typically found in promoters (e.g., SP/KLF, ETS factors, and USF). Multiple CREs are also regulated by NRF1, a TF involved in processes like metabolism, DNA repair, and cell cycle, all related to viral replication [54]. We also detected many CREs regulated by TFs acting downstream of immune and stress activation pathways, such as FOS:JUN, ATF2, CREB1, IRF3, and STAT [6]. These findings suggest that adenovirus CREs sense host activation pathways to activate transcriptional programs promoting viral replication.

In addition to adenovirus 5, we tested seven other adenoviruses from species B (types 3, 7, 11, and 14), C (types 1 and 5), D (type 37), and E (type 4). We detected activity conservation across strains for known promoters, with the L4/E2L promoter being the most variable and the MLP being the most conserved (**Figure 6C**). This activity conservation is likely partly related to conservation in regulation by SP, ELF, CREB1, and NFY factors (**Figure 6D**). However, we also detected marked variation in regulators both across and within species, particularly for TFs such as FOS:JUN, GFI1B, ATF2, and ZBTB33, suggesting varying responses across viral types.

The activity of most novel CREs is less conserved across adenovirus types. For example, CREs in the L1-L4 region show low activity conservation (**Figure 6C**). While these CREs are abundant and conserved between types 1 and 5 from species C, many are lost in species B types (3, 7, 11, and 14). Similarly, two CREs located between the UXP and E4 promoters are highly active and conserved between types 1 and 5, but one or both are lost or markedly reduced in other types. This low conservation is also observed across isolates from the same adenovirus type (**Figure S6A**). While the impact of the CREs on gene regulation is still unknown, these findings suggest extensive regulatory plasticity between and within viral types.

### CRE activity in adenoviral vectors

The neutral tropism and large genome capacity of adenoviruses have been harnessed to design viral vectors for gene therapy and vaccines, including the Vaxzevria, Janssen, Sputnik V, and Convidecia Covid-19 vaccines, and Zabdeno for Ebola [55,56]. Three generations of adenoviral vectors have been developed that delete different genomic regions to prevent the generation of replication competent viruses, avoid the expression of oncogenic proteins, and to produce space for transgenes. Given that these vectors retain parts of the adenovirus genome, they may contain CREs that can influence transgene expression.

Adenovirus 5 is often used as a backbone for these vectors, with all three generations retaining a ∼450 bp sequence containing the inverted terminal repeat and Ψ sequences, necessary for DNA replication and packaging. This region is highly active in MPRAs (30-200-fold) across cell lines (**Figure 6C**). This is important, as generation 1 and 2 vectors often place the transgene downstream of this element. Although this can be advantageous for constitutive transgene expression, this active Ψ CRE may lead to high background expression or unintended interactions with CREs controlling transgenes in an inducible or tissue-specific manner. *In silico* saturation mutagenesis using Malinois identified six ELF, one FOS:JUN, one HNF4A, one SP/KLF, and one USF sites in the Ψ region, which do not overlap with the packaging sequences (**Figures 6E** and **S6B**). We introduced mutations in these sites predicted to abrogate the transcriptional activity (**Figure 6F**). Next, we constructed Adenovirus 5 vectors containing the wild type or mutant Ψ sequences using the Adenobuilder kit, and evaluated the effect on viral production and activation of a doxycycline inducible promoter driving GFP expression. Mutant Ψ did not reduce viral titer, suggesting that DNA packaging was not affected **(Figure 6G**). The Ψ mutations reduced basal GFP levels by 65% in the uninduced condition, but only by 30% at saturating doxycycline concentration (**Figure 6H**), resulting in a 2-fold increase in inducibility in the mutant virus (**Figure 6I**). These findings show that adenoviral vectors can be optimized for inducibility by rationally mutating known or cryptic CREs, while preserving viral functions.

### Herpesviruses differ in CRE types and genomic distributions

Human Herpesvirus genomes range from 125 to 230 kb, encoding tens of genes. These viruses have complex regulatory mechanisms, following a highly regulated cascade, with immediate early genes initiating replication, late genes encoding structural proteins for virion assembly and egress, and latent genes essential for persistence [5]. These processes require multiple CREs and a tight interplay between host and viral TFs [5,6]. Indeed, we detected 89-268 CREs per virus per cell line (**Figure 1B** and **Table S4**) and 477-3,176 TF-CRE interactions per virus predicted by Malinois in K562 cells (**Table S6**), highlighting the regulatory complexity of herpesviruses.

The CRE repertoire varies across viruses. For example, most active tiles from HHV-1, HHV-2, HHV-5, and HHV-6B are promoter-like, while active tiles from other viruses are distributed across CRE classes, with HHV-7 being enriched in distal tiles (**Figures 2C** and **7A**). This does not necessarily imply that HHV-7 CREs function as distal elements, but rather that they are regulated by TFs frequently found in human distal elements (e.g., USF and IRF), are depleted in TFs found in promoter-like elements (e.g., SP and ELF), and have a similar dinucleotide composition to distal human CREs (**Figures 2H** and **S3I**).

**Figure 7:**
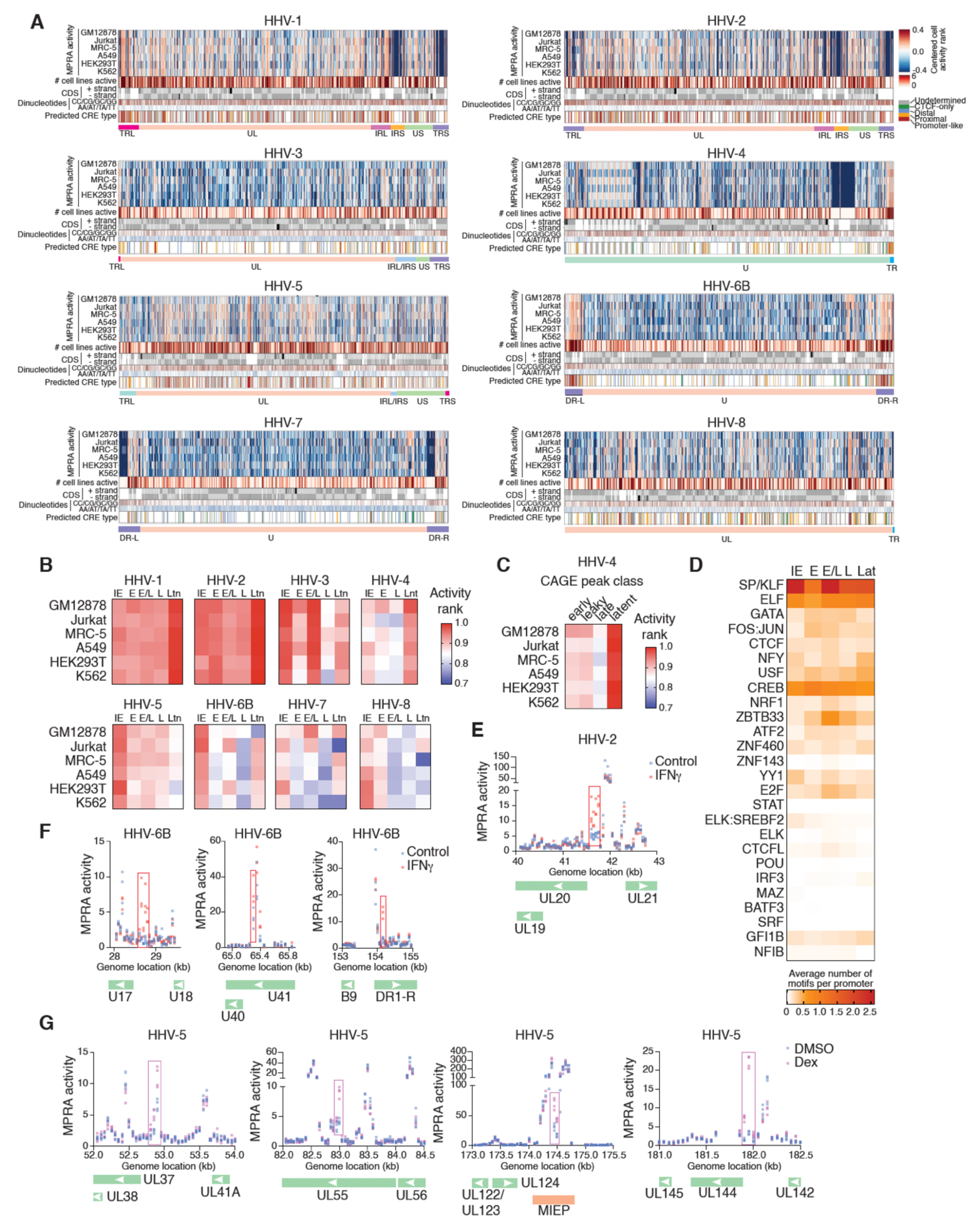
Landscape of Herpesvirus CREs. (A) CRE map across Herpesvirus species. MPRA activity is shown as a centered activity rank per cell line. The number of cell lines in which each tile was active is indicated. Coding sequences are shown, where dark gray indicates a coding sequence is in the corresponding strange, and light gray if it’s in the opposite strand. Dinucleotides frequencies are shown for CC/CG/GC/GG and AA/AT/TA/TT. For each active tile, the predicted CRE class is shown. The viral genome structures are shown at the bottom. (B) Median MPRA activity rank for gene promoters of immediate early (IE), early (E), early/late (E/L), late (L), and latent (Ltn) genes per virus per cell line. Promoters are defined as the regions ± 250 bp from the annotated 5’ end of genes. (C) Median MPRA activity rank for HHV-4 tiles that show CAGE peaks and were classified as early, leaky, late, and latent. (D) Average number of regulatory TF motifs for promoter regions corresponding to genes classified as immediate early, early, early/late, late, and latent across all Herpesviruses tested. Only TFs with an average of at least 0.1 motifs in at least one timepoint are shown. (E, F) Regions of HHV-2 (E) and HHV-6B (F) activated by incubation of K562 cells with 100 ng/ml of IFN𝛾 for 6 hs. (G) Regions of HHV-5 activated by incubation of K562 cells with 100 nM of dexamethasone for 6 hs.

CREs are distributed differently across viral genomes (**Figure 7A**). HHV-1, HHV-2, and HHV-5 have highly active CREs, evenly distributed across their genomes. Other herpesviruses show non-uniform CRE distributions. HHV-3 has fewer and less active CREs, with a higher density in the IRL/IRS, US, and TRL regions, enriched in genes transcribed in latently infected cells [57]. HHV-4 has a high density of promoter-like tiles in the terminal repeat region and tiles of multiple classes across the rest of the genome. HHV-6B and HHV-7 promoter-like tiles are highly active and concentrated in the direct repeats, while distal and CTCF-only tiles are sparse and have low activity. HHV-8 shows an even distribution across CRE classes, with promoter-like tiles being the most active. Altogether, this illustrates the complex spatial organization of CREs across herpesvirus genomes.

We found that promoter regions (± 250 bp of the annotated 5’ end) of genes expressed at different viral replication stages had varying activity levels (**Figure 7B**). For HHV-1, HHV-2 and HHV-4 latent gene promoters were the most active across cell lines. This is supported by analyzing tiles with CAGE peaks [19] at different stages during HHV-4 reactivation (**Figure 7C**). Conversely, for other herpesviruses, immediate early or early gene promoters were often the most active (**Figure 7B**). Late genes promoters had low activity, likely due to MPRAs being performed in non-infected cells. Indeed, about half of early and leaky HHV-4 TSSs reside within active tiles, while only 16% of late TSSs do so, suggesting that other viral or host factors drive late TSS activity during infection. Given the observed differences, we hypothesized that different host TFs would regulate immediate early, early, late, or latent genes. Indeed, SP and YY1 preferentially regulate immediate early genes, CREB and FOS:JUN regulate early genes, and NFY, USF, and GFI1B regulate latent genes (**Figure 7D**). However, these general trends are not conserved across all herpesviruses (**Figures 2J, S4B,** and **S7A**).

### MPRAs identify herpesvirus CREs that respond to cellular stimulation

Viral CREs can act as sensors to produce viral proteins that promote replication or avoid immune responses. Interferon gamma (IFN𝛾) plays a central role in antiviral responses by activating immune cells, inducing an antiviral state, enhancing antigen presentation, and promoting cell death [58,59]. To identify viral CREs activated by this cytokine, we performed MPRAs in K562 cells stimulated for 6 hours with 100 ng/ml of IFN𝛾. We detected an IFN𝛾-responsive region containing a STAT1 motif in HHV-2 upstream of UL19, UL20, and UL21, which encode for capsid, viral egress, and DNA packaging proteins that promote viral propagation (**Figure 7E**). Additionally, UL21 promotes cGAS degradation [60], blocking a key antiviral mechanism positively regulated by IFN𝛾. Interestingly, the homologous region in HHV-1, which lacks the STAT1 site, does not respond to IFN𝛾 (**Figures S7B-D**).

We also identified three HHV-6B CREs highly responsive to IFN𝛾: upstream of U17, upstream of U40, and downstream of the DR1-R/DR1-L TSSs, all containing STAT1 binding sites (**Figure 7F**). U17 and DR1-R are structural homologues of HHV-5 UL36, which interacts with caspase 8 to prevent cell death [61]. Since IFN𝛾 promotes caspase 8-mediated apoptosis of infected cells, this suggests HHV-6B may counteract this by increasing the production of anti-apoptotic proteins in response to IFN𝛾.

Many herpesviruses are reactivated by stress, such as high cortisol levels [62], and by glucocorticoid treatments, both due to immune suppression and activation of immediate early genes [62–65]. For example, the HHV-5 major immediate early promoter, which drives IE1/IE2 expression, is activated by dexamethasone through a glucocorticoid response element [63]. To identify other glucocorticoid-responsive HHV-5 CREs, we performed MPRAs in K562 cells stimulated with 100 nM dexamethasone for 6 hs. In addition to the major immediate early promoter, we found three other activated regions, all with glucocorticoid receptor binding sites, close to immediate early genes UL37, UL55, and UL144 (**Figures 7G** and **S7E**). These genes inhibit immune responses, block apoptosis, and promote viral replication and spread [66–69]. Altogether, this suggests that glucocorticoids contribute to HHV-5 reactivation and replication through upregulation of multiple immediate early genes.

## Discussion

Viral CREs have been identified and characterized through mechanistic studies of individual elements or genome-wide using histone marks, TF binding profiles, and sites of transcriptional initiation for a subset of viruses. This work greatly expands on previous studies, providing a comprehensive functional map of the CRE landscape for 27 dsDNA human-infecting viruses. Using MPRAs, we identified ∼2,000 CREs and observed marked differences in CREs across and within viral families, including variations in genomic CRE distributions, activity levels, and contributing TF motifs. This data is available as a user-friendly web resource (https://www.vcre-vis.org/) with visualizations of CRE activity across cell lines, activity filters, and motif predictions. Consistent with previous studies of individual elements [6] (**Table S1**), we found that viral CREs share regulatory features with mammalian CREs, being controlled by similar TF repertoires.

However, we also found key differences: first, viral genomes have higher CRE density, with an enrichment in promoter-like and a depletion of distal CREs; second, about half of viral CREs were classified as undetermined, sharing features of promoter-like or proximal elements, but being enriched in coding sequences and showing lower MPRA activity; third, most viral CREs overlap with coding sequences, which is rare for human CREs, potentially imposing reciprocal variation constraints and adding to constraints associated with overlapping viral genes [70].

Viral isolates from the same type can display extensive sequence variability, especially in families like Polyomavirus and Papillomavirus [71,72]. This variability has been linked to differences in regulation, resistance development, and pathogenesis. We show that some CREs, though mostly conserved at the sequence level, display extensive activity variability, which can sometimes be traced to few variants within motifs critical for CRE activity. This activity variability can provide a substrate for regulatory variation, response to different signaling pathways, and strain selection under selective pressure. For example, variation in NFAT binding sites in the NCCR of BK polyomavirus has been associated with differences in transcriptional activity, viral production, and reactivation in renal transplant patients [73].

Studies of Adenovirus 5 generation 1 vectors have shown that transgene expression levels depend on the promoter type and transgene orientation [74,75]. Our results suggest that this could be related to the numerous CREs in the adenoviral genome, which may contribute to transgene expression proximally or distally. The use of tunable promoters for transgene expression, desirable in some gene therapy and vaccine applications, may require removing or mutating CREs entirely. Our predicted motif contributions helped select mutations in the Ψ region to improve transgene inducibility, while preserving essential functions for viral production. Since the Ψ region was active across all adenoviruses tested, our approach could be used to increase inducibility in other adenoviral vectors. More generally, this design pipeline could be applied to adeno-associated, herpesvirus, and lentiviral vectors.

This study greatly expands the viral CRE landscape, but additional CREs may still be uncovered in different cell types or conditions, as some CREs are cell line specific. Further, the activity of many CREs may be modulated by cellular pathways that impinge on various host TFs. Previous studies have shown that many viral CREs are regulated by signal- or ligand-activated TFs, such as nuclear hormone receptors, NF-κB, TP53, and HIF1A (**Table S1**). We show that IFN𝛾 activates several HHV-2 and HHV-6B CREs, whereas dexamethasone activates HHV-5 CREs. Therefore, the 26 TF motifs we found using Malinois are likely an underestimate, given that Malinois and TFmodisco-lite may miss low frequent motifs, and we used Malinois trained in only one cell line in unstimulated conditions.

In addition to host TFs, many viral CREs are regulated by viral TFs. Other viral proteins can also modulate signaling pathways, interact with host TFs, hijack the host transcriptional machinery, reprogramming host gene regulatory networks that control viral CRE activity. For example, in HEK293T cells that expressed SV40 large T-antigen, we found that this viral TF contributes to JC polyomavirus NCCR activity. MPRAs in GM12878 cells, latently infected with HHV-4 revealed that latent gene promoters regulated by latent viral TFs such as EBNA2, EBNA1, EBNA-LP, and EBNA3, were among the most active HHV-4 CREs. Expanding this study to explore different conditions, infection states, and activation mechanisms of viral CREs will be key to identifying triggers of viral reactivation or pathogenic protein upregulation. These studies could inform precision sensitizing approaches to increase the expression of antigenic viral proteins, which, combined with vaccines, could help eliminate latently infected cells, similar to shock-and-kill strategies proposed for HIV [76,77]. Finally, our studies should eventually be integrated with target gene identification using perturbation or chromatin conformation studies as performed for HHV-4 and HHV-8, among other viruses [78,79].

## Methods

### Literature curation of viral CRE-TF interactions

To obtain a comprehensive list of TFs that bind or regulate different viral CREs, we manually searched PubMed using the virus name and combinations of terms such as “transcription factor”, “binding”, “regulation, “activation”, “repression”, “chromatin immunoprecipitation”, “reporter”, etc. The curation was performed by inspecting the full manuscript (rather than the abstract alone) by 2 independent researchers, one of them being a senior author. Publications that included the virus name, a viral CRE, a human TF, and experimental evidence for the interaction were included in **Table S1**. In addition, we annotated whether the interaction is activating or repressing and the PubMed ID of the publication. Overall, we generated a table containing 449 unique interactions between 147 viral CREs and 123 human TFs.

### MPRA viral tiling library sequence design

Viral sequences used for generating the MPRA tiling library are listed in **Table S3**, including virus family and strain, source (e.g., accession number), and sequence length. Each virus was tiled using 200 bp sequences with 50 bp offsets and each tile was tested in both orientations relative to the minimal promoter in the MPRA vector. Tile generation begins from the 5’ end of the sequence provided for each virus and the offset applies until the 3’ end of the sequence. To avoid having tiles shorter than 200 bp, the last tile for each virus corresponds to the last 200 bp sequence from the 3’ end. Sequences for adenovirus 1, 5, 7, 14, and 37 were obtained from ATCC and did not include some of 5’ and 3’ genomic sequences. To recover these sequences, we identified corresponding full-length genomes, matching serotype and sequence context, from NCBI and supplemented the missing 5’ and 3’ ends of the ATCC entries. This ensured full representation of both terminal regions for the adenovirus strains in the MPRA library. Overall, we generated 66,026 tile sequences for the virus listed.

In addition, we included negative and positive control sequences in the MPRA library to evaluate the sensitivity of each MPRA experiment, and to normalize activity for experimental tiles. Negative controls consisted of 250 shuffle sequences, 1,876 sequences overlapping human ORFs, and 506 sequences without detected MPRA activity. The positive control set of tiles consisted of 91 human genome sequences known to be active in MPRA, representing a broad activity spectrum. These control sequences correspond to those used in our previous studies [14].

### MPRA Library Construction

The MPRA libraries were constructed as previously described [80]. Briefly, oligonucleotide pools were synthesized (Twist Biosciences) as 230 nt sequences containing 200 nt sequences (or from 180-200 nt for the isolates) and 15 nt of adaptor sequence on both ends. Unique 20 bp barcodes were added by PCR (**Table S8**), along with additional constant sequences, for subsequent incorporation into a backbone vector (Addgene #109035) by Gibson assembly. The oligonucleotide library was electroporated into NEB 10-beta *E. coli*, and the resulting plasmid library was sequenced using Illumina 2 × 150 bp chemistry to determine oligo-barcode pairings. The library then underwent restriction digestion using AsiSI, and GFP with a minimal TATA promoter was inserted by Gibson assembly, resulting in the tile sequence being positioned directly upstream of the promoter and the 20 bp barcode residing in the 3′ UTR of GFP. After expansion in *E. coli*, the final MPRA plasmid library was sequenced using Illumina 1 × 26 bp or 2 x 150 bp chemistries to acquire a baseline representation of each oligo-barcode pair within the library.

Two separate libraries were generated using this method. The first library contained the viral genome tiling sequences described above as well as the negative and positive control tiles. The second library contained two sets of test sequences: saturation mutagenesis and viral isolate sequences of polyomavirus and papillomavirus CREs, in addition to the negative and positive controls.

### MPRA library transfection into cell lines

Jurkat cells (ATCC-TIB-15) and K562 cells (ATCC-CCL-243) were grown in RPMI media with 10% FBS, and GM12878 cells (Coriell) in RPMI + GlutaMAX supplemented with 15% FBS, up to density of 1 million cells per mL prior to transfection. HEK293T cells (ATCC-CRL-11268) were grown in DMEM with 10% FBS, MRC5 (ATCC-CCL-171) were grown in EMEM with 15% FBS, and A549 (ATCC-CCL-185) were grown in F12K media with 10% FBS. 3 electroporation replicates were conducted for A549, 4 replicates for K562, HEK293T, GM12878, and MRC-5, and 5 replicates for Jurkat cells. Each replicate was performed by electroporating 100 million cells with 100 ug of MPRA plasmid. Electroporation was performed using the 100uL Neon Transfection System kit with 3 pulses of 1100V for 10ms for HEK293T cells, 3 pulses of 1350V for 10 ms for Jurkat cells, 2 pulses of 1150V for 30ms for A549 cells, 2 pulses of 1200 V for 40ms for MRC-5, 3 pulses of 1450 V for 10ms for K562 cells, and 3 pulses of 1200V for 20ms. Cells were then incubated for 24 hs in the corresponding media to allow for expression of the barcoded reporter construct. Finally, cells were pelleted, washed three times with PBS, and stored at –80 °C for later RNA extraction.

### RNA extraction, GFP pull down, and MPRA RNA-seq library generation

RNA from all cell lines was extracted from frozen cell pellets using the Qiagen RNeasy Maxi kit. GFP transcripts were pulled down, retrotranscribed, and libraries were prepared for sequencing as previously described [14]. Briefly, half of the isolated total RNA underwent DNase treatment and a mixture of three GFP-specific biotinylated primers (#120, #123 and #126)(**Table S8**) were used to capture GFP transcripts with Streptavidin C1 Dynabeads (Life Technologies), followed by an additional DNase treatment. cDNA was synthesized from GFP mRNA using SuperScript III(Thermo Fisher Scientific) and purified with AMPure XP beads (Beckman Coulter). Quantitative PCR using primers specific for the GFP transcript (#781 and #782)**(Table S8**) was used to measure GFP transcript abundance in each sample. Replicates within each cell type were diluted to approximately the same concentration based on the qPCR results. Illumina sequencing libraries were constructed using a two-step amplification process to add sequencing adapters and indices. An initial PCR amplification with NEBNext Ultra II Q5 Master Mix and primers #781 and #782 was used to extend adapters. To minimize over-amplification during library construction, the number of PCR cycles used in the first amplification was selected based on where linear amplification began for each cell type. A second 6 cycle PCR using NEBNext Ultra II Q5 Master Mix added P7 and P5 indices and flow cell adapters (**Table S8**). The resulting MPRA RNA-tag libraries were sequenced using paired-end Illumina chemistry and dual index reads on either a NextSeq 1000 or NovaSeq 6000 instrument

### MPRA quality control and data analysis

To determine the quality of the MPRA libraries, we calculated the number of barcodes per sequence, the correlation between experimental replicates, and the activity of positive and negative control sequences. The oligo library was covered by a mean of 324 barcodes in the plasmid library, 277 barcodes in HEK293T, 242 in Jurkat, 134 in MRC-5, 212 in A549, 270 in K562, and 71 in GM12878 cells (**Figure S1A**). We also determined that 97.7% of oligos in the plasmid library, 97.4% in HEK293T, 97.2% in Jurkat, 95.8% in MRC-5, 97.0% in A549, 97.3% in K562, and 93.1% in GM12878 libraries were recovered with >10 barcodes, respectively. We observed high correlations of oligo counts across biological replicates for all cell lines (pearson correlation coefficient of 0.98–1) (**Figure S1B**). We also show that our MPRA experiments in all six cell lines clearly distinguish the activity of positive and negative controls (**Figure S1C**). These measures illustrate the high-quality of the MPRA experiments.

Data from the MPRA experiments was analyzed as previously described [80,81]. The sum of the barcode counts for each oligo were provided as input to DESeq2 (v.1.28.0) and replicates were median normalized followed by an additional normalization of the RNA samples to center the average RNA/DNA activity distribution of the negative control sequences over a log2 fold change of zero (**Table S9**). This normalization was performed independently for each cell line.

### MPRA saturation mutagenesis and isolate library design

For saturation mutagenesis, we selected 39 active tiles (**Table S10**). We primarily focused on the major CREs controlling gene expression in three polyomaviruses (BK, JC, and Merkel Cell) and two papillomaviruses (types 16 and 52). We selected the tiles with strong CRE activity from the NCCR regions of the polyomaviruses and from the LCR regions of the papillomaviruses. To minimize redundancy, we avoided selecting tiles that overlapped by more than 100 bp. In addition, we selected CREs located outside the NCCR/LCR regions, particularly those with strong activity, and for papillomaviruses, tiles that were located in similar genomic regions across the studied viruses. Saturation mutagenesis was performed from positions 6 to 195 of each selected 200 bp tile by introducing all possible single-nucleotide substitutions, wherein each nucleotide position was individually mutated to the other three possible bases, resulting in a total of 570 unique single-nucleotide variants per tile tested. The mean variant effect was represented in the height of the letter in the logo sequence. Motifs were manually identified in the resulting plots using Tomtom [24].

For the tiles indicated above, we have also evaluated homologous sequences corresponding to isolates from the same viral species. In June 2023, we retrieved all publicly available complete genome assemblies for each target virus from the NCBI Virus portal, applying the “complete” filter to exclude partial or draft records and recording the download date to ensure reproducibility. Using Python and Biopython’s local alignment mode, we sliced each sequence into 200-nt tiles (including their reverse complements) and aligned them against every reference genome with a scoring scheme of +2 for matches, 0 for mismatches, and gap penalties of –20 (opening) and –2 (extension). From each alignment we extracted the aligned length and the number of exact matches for both forward and reverse orientations, then merged these results into a unified dataset for downstream analysis.

Next, we imported the alignment outputs into R and performed a global realignment using Biostrings’ pairwiseAlignment with a nucleotide substitution matrix (match = 2, mismatch = –3), a gap opening penalty of –5, and a gap extension penalty of –2. We filtered out any alignments shorter than 180 nt or containing fewer than 100 exact matches. To guard against overly extended outliers, we calculated the 95th percentile of the remaining alignment lengths for each tile and set a dynamic maximum length threshold: tiles whose lengths exceeded this threshold (removing at most the top 5 % of extensions) were discarded, while the remainder were retained. Tiles identified as originating from reverse-strand alignments were reverse-complemented so that all sequences share a uniform 5′→3′ orientation. In a final pass, any sequence still longer than 200 nt was trimmed to its last 200 nt, its start position adjusted accordingly, and its exact-match count recalculated via one more global alignment. This procedure yields a final collection of 200 nt tiles that uniformly satisfy our stringent length and match criteria. This library also included the same set of positive and negative control sequences as the tiling library above.

### Identification of active viral tiles and CRE definition

We used a cell-specific threshold separating positive from negative control tiles at 1% FDR to call viral tiles active. In each cell line we considered the distributions of log2 (FC + 1) values for the negative control sequences and for the positive control sequences. For both positive and negative control distributions we trimmed the 2.5% extreme values from both ends. Then a threshold value of log2 (FC + 1) was chosen that separates the positive control from negative control tiles with the 1% FDR. Specifically, we have thus chosen activity threshold values of 1.634, 1.652, 1.801, 2.002, 2.114, 2.580 for the GM12878, Jurkat, MRC5, A549, HEK293, K562 cell lines, respectively. All viral tiles with activity greater or equal to this cell-specific threshold value were considered active. For some analysis, we aimed to compare tile activities across cell lines, in which case, we used the cell line activity ranks (indicated in the corresponding figures). To calculate these ranks, we ordered all tiles from all viruses by their activity in a corresponding cell line. To define cell-specific CRE positions, we used the union of positions of overlapping tiles active in at least one orientation. We note that this procedure is equivalent to applying bedtools [82] merge function to all active tiles in a given cell line.

### Cell line specificity of viral CREs

Since exact CRE positions may differ across cell lines, we re-defined CRE positions to calculate the number of shared CREs. For this purpose, we divided each CRE into subsegments which we call Partitioned CREs, such that each Partitioned CRE is either entirely active or entirely not active in each individual cell line. This was done using a custom Python function. We note that the procedure is equivalent to using the bedtools [82] intersect function for CREs of each cell line versus all other cell lines and then taking the union of all the resulting segments. To remove artifacts, we filtered out Partitioned CREs shorter than 200 bp. The resulting set of Partitioned CREs was used to quantify the number of shared CREs across cell lines per virus and to assess CRE size based on the number of cell lines in which a given CRE is active.

Additionally, we examined the cell line specificity of individual tiles. We used the supervised clustering approach where all the tiles were grouped into 64 classes based on the binary pattern of their activity in each of the 6 cell lines. Next all the classes were sorted by size and the 20 most frequent classes plotted in **Figure S2B**.

### Comparison between MPRA activity and experimentally identified or annotated TSSs

To determine the proximity between gene promoters and MPRA-identified CREs, we calculated a distance between each CRE midpoint (in HEK293T cells) to the nearest gene start coordinate available in the GenBank annotation (**Table S11**). Here, we refer to gene start coordinates as TSSs, though we note that the exact TSS positions may not correspond to gene starts provided in GenBank annotations. For each virus we calculated the fraction of CREs falling within the 250 bp proximity range of a nearest TSS. Given the density of TSSs and CREs in viral genomes, we expected these fractions to be high even if CRE positions were located at random. Thus, we compared the proximal CRE fractions to the expected values given the random uniform location of CRE center positions. The sample size for each virus was 10 times the number of CREs annotated in the corresponding virus.

Additionally, we did the reciprocal analysis where we calculated distances for each gene start position to the nearest CRE midpoint (in HEK293T cells). The fractions of TSSs being closer than 250 bp to the nearest CRE midpoint were compared to a random expectation given a uniform location of gene start positions. The expectation was estimated by sampling uniformly TSS positions across the viral genome with a sample size equal to 10 times the number of gene start positions provided in GenBank for the corresponding virus.

As an orthogonal approach, we measured the agreement between experimentally identified TSSs and MPRA activity. For this analysis, we used the data from Reza Djavadian et al. [19] for HHV-4 (V01555.2), István Prazsák et al. [20] for HHV-8 (GQ994935.1), Adam W. Whisnant et al. [12] for HHV-1 (BK012101.1), and Shirley E Braspenning et al. [18] for HHV-3 (NC_001348.1) (**Table S11**). Since the GenBank genome records for HHV1, HHV-3, and HHV-8 were different from the ones we used for other analysis, we re-mapped MPRA tile sequences to these genome records to retrieve the consistent TSS and tiles coordinates. We only considered the tiles aligning with no more than 5 bp mismatches. We then compared the distribution of activity of tiles containing a TSS to those without an identified TSS using the KS test. We also performed this analysis separately in coding and non-coding regions to control for the different TSS and MPRA activity distributions in these two genome region types.

### Motifs contributing to MPRA activity

To estimate how well the Malinois model [14] predicted MPRA activity for tiles obtained from viral genomes, we compared Malinois predictions and measured MPRA activity values in K562 cells across viral tiles. As the Malinois model was trained to predict an average between forward and reverse sequence orientation activities, we calculated the measured activity value for each tile as an average of log2 (FC + 1) values from both orientations. We then calculated the Pearson R correlations between predicted and experimental log2 (FC + 1) values across all tiles, and for tiles from individual viruses.

Motivated by the high correlation values of predicted and experimentally observed tile activities, we decided to utilize the Malinois model to identify TF motifs contributing to MPRA activity. We used the Sampled Integrated Gradients (SIG) method to calculate hypothetical contribution scores for the K562 cell line Malinois model as described previously (arXiv:1703.01365, 39443793). We calculated SIG values for all viral tiles active in K562 (totaling to 38778 sequences) as well as for a set of 157,766 human DHS sequences identified in [83].

The DHS positions were downloaded from https://resources.altius.org/~jvierstra/projects/footprinting.2020/per.dataset/ as a bed file (K562-DS15363). For each K562 DHS we considered the central 200 bp, corresponding to the size of the MPRA tiles that we used and to the input of the Malinois model. We considered DHS sequences together with viral CRE tiles to get a larger set of contribution score arrays and, thus, to increase the probability of detecting low-frequency motifs. We then applied TF-MoDISco-lite (arXiv:1811.00416, https://github.com/jmschrei/tfmodisco-lite (2022)) with 1000000 seqlets and a window size of 200 to extract sequence motifs frequently occurring in SIG profiles. The resulting 47 motif matrices were compared against the SCENIC motif collection [84] using TOMTOM [24]. We also manually inspected motif matrices and assigned a TF group / family to each matrix. Single and composite TF motifs were collapsed into 26 single TF motifs. A motif resembling 3’ splice sites was excluded from further analysis as it is likely associated with transcription from a cryptic upstream promoter [85].

### TF motif annotations based on Malinois predictions

To assign TF motifs to each tile and determine their position, we scanned SIG arrays with the TF-MoDISco-lite identified motif matrices using the Continuous Jaccard Similarity (CJS) measure proposed in [86]. Briefly, for each tile we calculated the sum of absolute SIG values for all nucleotides at each position (total absolute SIG profile). We then assigned the contributing TF binding sites by applying a simple peak calling algorithm to the resulting total absolute SIG profile (**Table S6**). This was done by calculating sliding window sums, with a window size of 5 bp, and then determining the maximums of these sums with the non-maximum suppression correction. To assign a TF-MoDISco-lite pattern (TF matrix) to each motif instance, we calculated the CJS for each TF matrix and the SIG matrix at a motif instance site and the assigned the TF matrix having the best CJS value to each motif peak.

### CRE classifier model architecture

We aimed to build a deep learning model predicting ENCODE CRE types for 200 bp DNA sequences active in the K562 MPRA. We reasoned that a model based on the pre-trained Malinois model might work well for this task. We made this assumption based on two observations: (1) the Malinois model has learned motifs of TFs contributing to either distal enhancer (Jun/Fos, GATA) or promoter-like (SP, ETS, CREB) activity; (2) the activity values predicted by Malinois for ENCODE Promoter-like elements were on average significantly higher than those predicted for ENCODE Distal elements, naturally providing a separation between these two CRE types.

Thus, we utilized the Malinois model backbone to build a CRE classifier convolutional neural network (CRE-class). Specifically, we completely transferred the convolutional part of the Malinois model architecture, consisting of three convolutional layers with batch normalization and maximum value pooling [14]. The weights of these three layers were initialized with those of the pre-trained Malinois model. We further added two fully connected layers serving as a classification head of the CRE-class model. The first linear layer projects the outputs of the convolutional part after flattening (vectors of size 2600) into 100 internal features, while the second layer further projects 100 internal features into 8 logit values for each of the target classes. Since the CRE-class convolutional layers correspond to those of the Malinois model, the input size to CRE-class is the same as for the Malinois, i.e., 600 bp one-hot encoded DNA sequences with the constant 200 bp flanks on each side.

### Data preprocessing and CRE classifier model training

We utilized a subset of ENCODE CREs (32728249) as the training dataset for the CRE-classifier model. For each ENCODE CRE we collected the central 200 bp region as a representative sequence of this CRE. After that we applied the Malinois model to each CRE sequence and only considered CREs predicted by Malinois to be active in at least one of K562, HepG2, and SK-N-SH cell lines. Here we used the same activity thresholds as the ones used to define active viral tiles. In this way, the training set sequences have similar activity properties to the sequences to which we eventually applied CRE-class (i.e., 200 bp viral tile sequences active in K562 MPRAs). We then randomly split sequences of each ENCODE class in a 9 to 1 proportion for the training and validation steps, respectively (**Table S12**). After the splitting we considered each sequence in both forward and reverse orientations, since by construction ENCODE sequences are not strand-specific. During the training, we did sequence augmentation by shifting sequences randomly by ±50 bp in their native genome context. Furthermore, for each sequence class we constructed a paired shuffled class consisting of sequences with the preserved tri-nucleotide frequencies using the ushuffle tool (https://github.com/guma44/ushuffle).

We trained a model to perform a classification into 8 classes using a label smoothing cross entropy loss. During the training we froze weights of the first two convolutional layers of the Malinois model, while the weights of the third layer were not freezed. To measure the model performance, we calculated the recall and precision for each class on the validation set. To confirm that the classifier model is making decisions based on both dinucleotide and TF motif sequence composition, we calculated the contribution scores (Sampled Integrated Gradients) for the train and validation sequences for the predicted logits for each of the output classes as described previously [14,87] and manually inspected the corresponding logos.

### CRE classifier inference for viral tiles

We applied CRE-class to viral tiles active in the K562 cell line. We note that the CRE-classifier model was trained to be not specific to sequence orientation (forward / reverse strand). Thus, to assign a class to a tile sequence we applied the model to both forward and reverse sequence orientations. As a result, we got 8 logits for each class for both orientations (**Table S5**). We further averaged logit values for each class between two sequence orientations. The class with the maximum assigned forward-reverse averaged logit value was considered to be the model’s prediction. Since we were interested in identifying viral tiles similar to real human CREs, we treated all tiles predicted as one of 4 shuffled classes as belonging to a single ‘Undetermined’ class. However, we note that in the vast majority of ‘Undetermined’ tiles corresponded to the Promoter-like shuffled class.

### Dinucleotide composition of human and viral tiles

We calculated the frequencies of each of the 16 dinucleotides for each tile from a set of ENCODE elements and viral tiles. In the case of ENCODE elements, we used the same set used for the CRE classifier model training and validation (see above). Each tile was converted to a vector of size 16 storing dinucleotide frequencies. Then, we performed Principal Component Analysis (PCA) on these vectors. For visualization purposes, we computed the Gaussian Kernel density estimation of the dot density on the PC-1 - PC-2 plane, where each dot corresponds to a single tile. Density estimation was performed separately for each tile group: ENCODE Promoter-like, ENCODE distal, and viral tiles. We additionally calculated average dinucleotide frequencies of tiles grouped by individual viruses and by predicted CRE types.

### Classification and characterization of viral tiles in coding and non-coding regions

To inspect the extent to which viral CREs overlap with coding regions (CDSs), we used CDS positions annotated by GenBank and assigned a CRE to be located in CDS if the midpoint of the CRE is within a CDS (**Table S5**). CDSs cover a substantial proportion of viral genomes and thus, expectedly, most CREs tend to be located in CDSs. We compared the observed fractions of CREs in CDSs with the expected fractions under the null model of CRE positions being distributed randomly uniform across the genome. The expected fraction of CREs is therefore equal to the genome fraction covered by CDSs.

To determine differences between active tiles in CDSs and non-coding regions, we calculated the distributions of activities in difference cell types, the density of TF motifs in the corresponding tile set, the enrichment in TSS in active versus inactive tiles, the proportion of tiles from is class (i.e., promoter-like, proximal, distal, CTCF-only, and undetermined), and the activity of tiles containing experimentally determined TSSs (**Table S11**).

### Assessment of sequence and MPRA-activity conservation across adenovirus isolate genomes

To evaluate sequence and activity conservation across adenovirus isolates from different types, we retrieved all available isolate genome sequences for the studied adenovirus types from GenBank on May 2025. For each type, we performed multiple genome alignments using the Pangenome Cactus tool [88]. Analyses were centered on the genome used for the MPRA screen, hereafter referred to as the MPRA-reference genome. Using the halLiftover tool [89], we mapped each MPRA-reference genome tile to its corresponding coordinates in the aligned isolate genomes. The midpoint between the start and end positions was used to define a 200 bp tile for each isolate (i.e., 100 bp upstream and downstream of the midpoint). Sequence identity between each isolate tile and the corresponding reference tile was computed using Biopython [90]. Predicted MPRA activity in K562 cells for each isolate tile was obtained using the Malinois model. This approach enabled us to quantify both sequence conservation and predicted regulatory activity across isolates for each tile in the MPRA-reference genome.

### Overlap between TF motifs and protein secondary structure features

To assess the overlap between TF motifs predicted to contribute to MPRA activity in K562 cells and protein secondary structure features, we used the Viro3D database [91], which provides AlphaFold-predicted protein structures for all viral genomes studied, with the exception of adenovirus type 4. The genome record used for protein structure prediction in adenovirus 4 was not included among the sequences analyzed in this study. Secondary structure annotations were generated using the DSSP algorithm [92,93] applied to AlphaFold output files. Amino acid positions were then mapped back to their corresponding genomic codons, allowing the assignment of a DSSP-derived secondary structure label to each codon. These labels were binarized into either ‘intrinsically disordered regions’ (IDRs) – corresponding to DSSP’s ‘Coil / None’ category – or ‘structured regions’, comprising all other DSSP-defined secondary structure types. We intersected codon-level secondary structure annotations with TF motif instances to quantify the number of codons overlapping each TF motif - structure category combination. We also computed the total number of codons encompassed by TF binding sites.

### Cloning of wild type and mutant **Ψ** constructs and adenovirus 5 vector assembly

We used Malinois trained in K562, SK-N-SH, and HepG2 cells to predict TF binding motifs within the Ψ region of Adenovirus 5 that contribute to transcriptional activity. We then introduced 1-3 mutations per motif at high activity contributing bases, while preserving known packaging elements. The resulting mutated sequence was re-evaluated by Malinois to confirm a reduction in predicted activity. Both wild-type (WT) and mutant Ψ sequences were then synthesized by Twist Biosciences. Full sequences of WT and mutant Ψ constructs, as well as primer sequences used for PCR amplification in the subsequent cloning steps, are listed in **Table S13**.

A Tet-responsive promoter fragment — comprising a Tet operator upstream of a minimal CMV promoter driving GFP (5′ to 3′ orientation) — was PCR-amplified from Addgene plasmid #115495 using primers Tet-ON-GFP-F and Tet-ON-GFP-R. To prepare the recipient vector, the Block 1 ΔE1-GFP plasmid (Addgene #179203), part of the Adenobuilder toolkit [94], was digested with XhoI and Bsu36I to excise the original Psi-CMV–GFP cassette, leaving the vector backbone intact.

The synthesized WT or mutant Ψ fragments and the Tet–CMV–GFP PCR product were assembled into the digested Block 1 ΔE1-GFP plasmid backbone using Gibson Assembly (NEBuilder HiFi DNA Assembly Master Mix, NEB Cat. #M5520A) in a 10 µL reaction at 50°C for 1 hour. Gibson products were transformed into chemically competent E. coli DH5α, and colony PCR was performed using primers Ad5-F and Tet-ON-GFP-R to verify insert presence and size. Correctly assembled plasmids were then confirmed by whole-plasmid sequencing using Plasmidsaurus and Azenta.

### Generation of recombinant adenovirus 5 vector, titration, and GFP activity measurement

The full-length recombinant Adenovirus 5 genome was assembled by Gibson Assembly using our modified Block 1 ΔE1-GFP construct and Blocks 2–7, comprising different overlapping segments of the viral genome, provided in the Adenobuilder toolkit [94]. The assembled genome was transfected into HEK293 Tet-On® 3G cells (Takara Cat. #631185) using Lipofectamine 3000 (Invitrogen Cat. #L3000-015), following the manufacturer’s instructions. Cells were maintained in DMEM supplemented with 10% Tet-system approved FBS (Takara Cat. #631105), 100 μg/mL

G418 (GoldBio Cat. #G-418-1), and Antibiotic-Antimycotic (Thermo Fisher Cat. #15240062). Viral harvest and amplification were done as described in the Adenobuilder protocol [94].

To determine viral titers, 1 × 10⁵ cells were seeded in 24-well plates and incubated for 48 hours prior to infection. Cells were then infected with different volumes of the virus preparation, along with 100 ng/mL doxycycline in the culture media, and incubated for 16 hours. GFP-positive cells were quantified by flow cytometry, and Poisson-based MOI estimation was used to calculate the multiplicity of infection based on the percentage of GFP-expressing cells.

To quantify GFP expression driven by the wt and mutant Ψ region, 1 × 10⁵ cells were seeded in 24-well plates and incubated for 48 hours prior to infection. Cells were then infected with the WT or mutant virus at an MOI of 5, along with 0.1-100 ng/mL doxycycline in the culture media, and incubated for 16 hours. Median GFP fluorescence was quantified by flow cytometry using a CytoFlex LX.

### Activity of herpesvirus gene promoters based on expression timing

Gene kinetic classifications for herpesviruses were primarily based on Fields Virology [5]. When annotations were outdated or missing, we supplemented them with peer-reviewed literature provided in **Table S14**. For consistency, specific temporal annotations such as ‘delayed early,’ ‘leaky late,’ and other subclassifications were consolidated into five broad categories: immediate early, early, early-late, late, or latent. For HHV-8, classifications were based solely on a publication [95]. We next collected gene start coordinates available at the GenBank annotations and refer to them as gene 5’ ends, and defined gene promoter regions as the region >250 bp away from the 5’ end. We then calculated the maximum cell MPRA activity rank between all tiles whose midpoint resides within this promoter region and assigned that activity rank value to a promoter. Finally, we determined the average promoter activity rank values across all genes within a kinetic group, and plotted this value for each herpesvirus and each cell line. In addition, we calculated the frequency of each TF motif derived from Malinois within the promoters of each kinetic group.

Since gene start positions provided in GenBank annotations do not necessarily correspond to actual gene promoters, we also analyzed an HHV-4 CAGE dataset (29864140) that defined TSS positions and their kinetic classes. In this study the authors adhered to a somewhat different gene kinetic classification with groups being: Early, Leaky, Late, Latent. The Leaky group in this classification corresponds to genes previously classified as Late, but found out to have a combination of Early and Late kinetic properties as defined by the expression pattern and dependence on the viral replication machinery. Therefore, in this analysis we used the kinetic classification from the original paper when calculating MPRA activity for HHV-4 tiles containing TSSs.

### MPRA in K562 cells stimulated with IFNG and dexamethasone

K562 cells (ATCC-CCL-243) were cultured in RPMI media with 10% FBS and 1% antibiotic antimycotic to a density of 1 million cells/mL prior to transfection. Four independent electroporation replicates were conducted, each performed by collecting 50 million cells and electroporating them with 25 ug of MPRA plasmid library. Cells were electroporated as previously described, and left to recover at 37 °C. After 24 hours, cells were treated with 100 nM dexamethasone or 100 ng/mL IFNg and their vehicle controls (0.06% DMSO and water, respectively). Following a 6 hour incubation, cells were pelleted, washed with PBS, and stored at –80 °C. RNA extraction and MPRA RNA-seq library generation were conducted as described above. Analysis of differentially active regions was performed using the DEseq2 (v1.46.0) in R (v4.4.2), as part of the Bioconductor package. Raw DNA and RNA count data were used as input and a paired design was implemented to account for matched samples. Normalization, dispersion estimation, and differential testing were performed following the standard DESeq2 parameters. Tiles were considered as differentially expressed if the *p*-value was <0.01, and the absolute value of log2FoldChange was more than 1 (**Tables S15** and **S16**).

### Resource Availability Lead Contact

Further information and request regarding the resources used in the manuscript should be directed to the lead contact, Juan Fuxman Bass.

### Materials availability

All the reagents generated in the study are available from the lead contact with a completed materials transfer agreement.

### Data and code availability

Code and scripts can be accessed at GitHub: https://github.com/osyafinkelberg/dsdna-mpra/tree/main/

## Acknowledgements

We thank Dr. Todd Blute for assistance with flow cytometry, Yun Shen for IT assistance installing and adapting packages in the computer cluster, and Dr. Leah Kottyan for comments on the manuscript. This work was supported by National Institutes of Health grants R35GM128625 (to J.I.F.B.), and R35HG011329 (to R.T.). J.A.F. was supported by NIH grant T32GM150533. E.M. was supported by the National Science Foundation grant BIO-1659605. J.P. was supported by Boston University’s undergraduate research opportunities program. L.M.C. was supported by ‘Programa de Prácticas de formación académica y profesional en el exterior para argentinos/a-RAICES-MINCYT’, Argentina.

## Author Contributions

Conceptualization, T.H.T, J.A.F., J.I.F.B.; methodology, T.H.T, S.K., B.E., B.D., E.M., J.P., J.R., M.P.; software, J.A.F., L.S-U., G.M-E.; formal analysis, J.A.F., T.H.T., L.S-U., B.E., H.C., R.C., G.M-E., R.T., J.I.F.B.; investigation, T.H.T., S.K., B.E., B.D., E.M., J.P., J.R.; resources, R.T. and J.I.F.B.; data curation, T.H.T., J.A.F., J.R., L.M-C., M.A.P., J.I.F.B; writing – original draft, T.H.T., J.A.F., J.I.F.B.; writing – review and editing, R.T. and J.I.F.B.; visualization, J.A.F., R.C., H.C., B.E., R.T., J.I.F.B.; supervision, R.T. and J.I.F.B.; project administration, J.I.F.B.; and funding acquisition, R.T. and J.I.F.B.

## References

1. Zur Hausen, H. (2001) Oncogenic DNA viruses. Oncogene 20, 7820–7823

2. Bjornevik, K. et al. (2022) Longitudinal analysis reveals high prevalence of Epstein-Barr virus associated with multiple sclerosis. Science 375, 296–301

3. Eyting, M. et al. (2025) A natural experiment on the effect of herpes zoster vaccination on dementia. Nature 641, 438–446

4. ElAbd, H. et al. (2025) T and B cell responses against Epstein-Barr virus in primary sclerosing cholangitis. Nat Med DOI: 10.1038/s41591-025-03692-w

5. Knipe, D.M. and Howley, P.M., eds. (2013) Fields virology, (6th ed.), Wolters Kluwer/Lippincott Williams & Wilkins Health

6. Rottenberg, J.T. et al. (2024) Viral cis-regulatory elements as sensors of cellular states and environmental cues. Trends Genet 40, 772–783

7. Kropp, K.A. et al. (2014) Viral enhancer mimicry of host innate-immune promoters. PLoS Pathog 10, e1003804

8. Davis, D.A. et al. (2001) Hypoxia induces lytic replication of Kaposi sarcoma-associated herpesvirus. Blood 97, 3244–3250

9. Haque, M. et al. (2003) Kaposi’s sarcoma-associated herpesvirus (human herpesvirus 8) contains hypoxia response elements: relevance to lytic induction by hypoxia. J Virol 77, 6761– 6768

10. Ostler, J.B. et al. (2019) The Glucocorticoid Receptor (GR) Stimulates Herpes Simplex Virus 1 Productive Infection, in Part Because the Infected Cell Protein 0 (ICP0) Promoter Is Cooperatively Transactivated by the GR and Krüppel-Like Transcription Factor 15. J Virol 93, e02063–18

11. Board, N.L. et al. (2022) Engaging innate immunity in HIV-1 cure strategies. Nat Rev Immunol 22, 499–512

12. Whisnant, A.W. et al. (2020) Integrative functional genomics decodes herpes simplex virus 1. Nat Commun 11, 2038

13. Fülöp, Á. et al. (2022) Integrative profiling of Epstein-Barr virus transcriptome using a multiplatform approach. Virol J 19, 7

14. Gosai, S.J. et al. (2024) Machine-guided design of cell-type-targeting cis-regulatory elements. Nature 634, 1211–1220

15. Marchini, A. et al. (2001) Human cytomegalovirus with IE-2 (UL122) deleted fails to express early lytic genes. J Virol 75, 1870–1878

16. Wiebusch, L. and Hagemeier, C. (1999) Human cytomegalovirus 86-kilodalton IE2 protein blocks cell cycle progression in G(1). J Virol 73, 9274–9283

17. Tai-Schmiedel, J. et al. (2020) Human cytomegalovirus long noncoding RNA4.9 regulates viral DNA replication. PLoS Pathog 16, e1008390

18. Braspenning, S.E. et al. (2020) Decoding the Architecture of the Varicella-Zoster Virus Transcriptome. mBio 11, e01568–20

19. Djavadian, R. et al. (2018) CAGE-seq analysis of Epstein-Barr virus lytic gene transcription: 3 kinetic classes from 2 mechanisms. PLoS Pathog 14, e1007114

20. Prazsák, I., et al. (2024) KSHV 3.0: a state-of-the-art annotation of the Kaposi’s sarcoma-associated herpesvirus transcriptome using cross-platform sequencing. mSystems 9, e0100723

21. ENCODE Project Consortium et al. (2020) Expanded encyclopaedias of DNA elements in the human and mouse genomes. Nature 583, 699–710

22. Nguyen, T.A. et al. (2016) High-throughput functional comparison of promoter and enhancer activities. Genome Res 26, 1023–1033

23. Gerstein, M.B. et al. (2012) Architecture of the human regulatory network derived from ENCODE data. Nature 489, 91–100

24. Gupta, S. et al. (2007) Quantifying similarity between motifs. Genome Biol 8, R24

25. Yordy, J.S. and Muise-Helmericks, R.C. (2000) Signal transduction and the Ets family of transcription factors. Oncogene 19, 6503–6513

26. Cheng, Z. et al. (2020) The interactions between cGAS-STING pathway and pathogens. Signal Transduct Target Ther 5, 91

27. Suzich, J.B. and Cliffe, A.R. (2018) Strength in diversity: Understanding the pathways to herpes simplex virus reactivation. Virology 522, 81–91

28. Feigenbaum, L. et al. (1992) JC virus and simian virus 40 enhancers and transforming proteins: role in determining tissue specificity and pathogenicity in transgenic mice. J Virol 66, 1176–1182

29. Moens, U. et al. (1995) Noncoding control region of naturally occurring BK virus variants: sequence comparison and functional analysis. Virus Genes 10, 261–275

30. Mitchell, P.J. et al. (1987) Positive and negative regulation of transcription in vitro: enhancer-binding protein AP-2 is inhibited by SV40 T antigen. Cell 50, 847–861

31. Romagnoli, L. et al. (2008) Early growth response-1 protein is induced by JC virus infection and binds and regulates the JC virus promoter. Virology 375, 331–341

32. Gorrill, T.S. and Khalili, K. (2005) Cooperative interaction of p65 and C/EBPbeta modulates transcription of BKV early promoter. Virology 335, 1–9

33. Yang, J.F. and You, J. (2020) Regulation of Polyomavirus Transcription by Viral and Cellular Factors. Viruses 12, 1072

34. Hashida, Y. et al. (2018) Genetic Variability of the Noncoding Control Region of Cutaneous Merkel Cell Polyomavirus: Identification of Geographically Related Genotypes. J Infect Dis 217, 1601–1611

35. Gosert, R. et al. (2008) Polyomavirus BK with rearranged noncoding control region emerge in vivo in renal transplant patients and increase viral replication and cytopathology. J Exp Med 205, 841–852

36. Kumar, M.A. et al. (2019) Directed Nucleosome Sliding during the Formation of the Simian Virus 40 Particle Exposes DNA Sequences Required for Early Transcription. J Virol 93, e01678–18

37. Lo Cigno, I., et al. (2024) High-risk HPV oncoproteins E6 and E7 and their interplay with the innate immune response: Uncovering mechanisms of immune evasion and therapeutic prospects. J Med Virol 96, e29685

38. McBride, A.A. (2022) Human papillomaviruses: diversity, infection and host interactions. Nat Rev Microbiol 20, 95–108

39. Morgan, I.M. (2025) The functions of papillomavirus E2 proteins. Virology 603, 110387

40. Taguchi, A. et al. (2020) Use of Cap Analysis Gene Expression to detect human papillomavirus promoter activity patterns at different disease stages. Sci Rep 10, 17991

41. Isok-Paas, H. et al. (2015) The transcription map of HPV11 in U2OS cells adequately reflects the initial and stable replication phases of the viral genome. Virol J 12, 59

42. Taguchi, A. et al. (2015) Characterization of novel transcripts of human papillomavirus type 16 using cap analysis gene expression technology. J Virol 89, 2448–2452

43. Milligan, S.G. et al. (2007) Analysis of novel human papillomavirus type 16 late mRNAs in differentiated W12 cervical epithelial cells. Virology 360, 172–181

44. Sawadogo, M. and Roeder, R.G. (1985) Interaction of a gene-specific transcription factor with the adenovirus major late promoter upstream of the TATA box region. Cell 43, 165–175

45. Wu, L. et al. (1987) A TATA box implicated in E1A transcriptional activation of a simple adenovirus 2 promoter. Nature 326, 512–515

46. Miller, D.L. et al. (2007) Adenovirus type 5 exerts genome-wide control over cellular programs governing proliferation, quiescence, and survival. Genome Biol 8, R58

47. Xie, D. et al. (2023) Oncolytic adenoviruses expressing checkpoint inhibitors for cancer therapy. Signal Transduct Target Ther 8, 436

48. Travieso, T., et al. (2022) The use of viral vectors in vaccine development. NPJ Vaccines 7, 75

49. Schwartz, U. et al. (2023) Changes in adenoviral chromatin organization precede early gene activation upon infection. EMBO J 42, e114162

50. Hsu, E. et al. (2018) Adenovirus E1A Activation Domain Regulates H3 Acetylation Affecting Varied Steps in Transcription at Different Viral Promoters. J Virol 92, e00805–18

51. Parks, C.L. and Shenk, T. (1997) Activation of the adenovirus major late promoter by transcription factors MAZ and Sp1. J Virol 71, 9600–9607

52. Fax, P. et al. (2000) cAMP-independent activation of the adenovirus type 12 E2 promoter correlates with the recruitment of CREB-1/ATF-1, E1A(12S), and CBP to the E2-CRE. J Biol Chem 275, 8911–8920

53. Kornuc, M. et al. (1990) Adenovirus early region 3 promoter regulation by E1A/E1B is independent of alterations in DNA binding and gene activation of CREB/ATF and AP1. J Virol 64, 2004–2013

54. Han, W. et al. (2012) Nrf1 CNC-bZIP protein promotes cell survival and nucleotide excision repair through maintaining glutathione homeostasis. J Biol Chem 287, 18788–18795

55. Jacob-Dolan, C. and Barouch, D.H. (2022) COVID-19 Vaccines: Adenoviral Vectors. Annu Rev Med 73, 41–54

56. Bulcha, J.T. et al. (2021) Viral vector platforms within the gene therapy landscape. Signal Transduct Target Ther 6, 53

57. Depledge, D.P. et al. (2018) A spliced latency-associated VZV transcript maps antisense to the viral transactivator gene 61. Nat Commun 9, 1167

58. Kak, G. et al. (2018) Interferon-gamma (IFN-γ): Exploring its implications in infectious diseases. Biomol Concepts 9, 64–79

59. Han, J. et al. (2023) Dysregulation in IFN-γ signaling and response: the barricade to tumor immunotherapy. Front Immunol 14, 1190333

60. Ma, Z. et al. (2023) Tegument protein UL21 of alpha-herpesvirus inhibits the innate immunity by triggering CGAS degradation through TOLLIP-mediated selective autophagy. Autophagy 19, 1512–1532

61. Chaudhry, M.Z. et al. (2020) Cytomegalovirus inhibition of extrinsic apoptosis determines fitness and resistance to cytotoxic CD8 T cells. Proc Natl Acad Sci U S A 117, 12961–12968

62. Glaser, R. and Kiecolt-Glaser, J.K. (2005) Stress-induced immune dysfunction: implications for health. Nat Rev Immunol 5, 243–251

63. Inoue-Toyoda, M. et al. (2015) Glucocorticoids facilitate the transcription from the human cytomegalovirus major immediate early promoter in glucocorticoid receptor- and nuclear factor-I-like protein-dependent manner. Biochem Biophys Res Commun 458, 180–185

64. Van Damme, E., et al. (2015) Glucocorticosteroids trigger reactivation of human cytomegalovirus from latently infected myeloid cells and increase the risk for HCMV infection in D+R+ liver transplant patients. J Gen Virol 96, 131–143

65. Yang, E.V. et al. (2010) Glucocorticoids activate Epstein Barr virus lytic replication through the upregulation of immediate early BZLF1 gene expression. Brain Behav Immun 24, 1089–1096

66. Ren, Y. et al. (2022) Dual inhibition of innate immunity and apoptosis by human cytomegalovirus protein UL37x1 enables efficient virus replication. Nat Microbiol 7, 1041–1053

67. Poole, E. et al. (2006) The UL144 gene product of human cytomegalovirus activates NFkappaB via a TRAF6-dependent mechanism. EMBO J 25, 4390–4399

68. Bitra, A. et al. (2019) Structure of human cytomegalovirus UL144, an HVEM orthologue, bound to the B and T cell lymphocyte attenuator. J Biol Chem 294, 10519–10529

69. Isaacson, M.K. and Compton, T. (2009) Human cytomegalovirus glycoprotein B is required for virus entry and cell-to-cell spread but not for virion attachment, assembly, or egress. J Virol 83, 3891–3903

70. Ho, J.S.Y. et al. (2021) Unconventional viral gene expression mechanisms as therapeutic targets. Nature 593, 362–371

71. White, M.K. et al. (2009) Regulation of gene expression in primate polyomaviruses. J Virol 83, 10846–10856

72. Burk, R.D. et al. (2013) Human papillomavirus genome variants. Virology 445, 232–243

73. Li, Y.J. et al. (2012) Cyclophilin A and nuclear factor of activated T cells are essential in cyclosporine-mediated suppression of polyomavirus BK replication. Am J Transplant 12, 2348– 2362

74. Suzuki, M. et al. (2015) Preferable sites and orientations of transgene inserted in the adenovirus vector genome: The E3 site may be unfavorable for transgene position. Gene Ther 22, 421–429

75. Farrera-Sal, M. et al. (2020) Effect of Transgene Location, Transcriptional Control Elements and Transgene Features in Armed Oncolytic Adenoviruses. Cancers (Basel) 12, 1034

76. Nixon, C.C. et al. (2020) Systemic HIV and SIV latency reversal via non-canonical NF-κB signalling in vivo. Nature 578, 160–165

77. Archin, N.M. et al. (2012) Administration of vorinostat disrupts HIV-1 latency in patients on antiretroviral therapy. Nature 487, 482–485

78. Morgens, D.W. et al. (2025) Enhancers and genome conformation provide complex transcriptional control of a herpesviral gene. Mol Syst Biol 21, 30–58

79. Morgan, S.M. et al. (2022) The three-dimensional structure of Epstein-Barr virus genome varies by latency type and is regulated by PARP1 enzymatic activity. Nat Commun 13, 187

80. Tewhey, R. et al. (2016) Direct Identification of Hundreds of Expression-Modulating Variants using a Multiplexed Reporter Assay. Cell 165, 1519–1529

81. Siraj, L., et al. (2024) Functional dissection of complex and molecular trait variants at single nucleotide resolution. bioRxiv DOI: 10.1101/2024.05.05.592437

82. Quinlan, A.R. and Hall, I.M. (2010) BEDTools: a flexible suite of utilities for comparing genomic features. Bioinformatics 26, 841–842

83. Vierstra, J. et al. (2020) Global reference mapping of human transcription factor footprints. Nature 583, 729–736

84. Bravo González-Blas, C., et al. (2023) SCENIC+: single-cell multiomic inference of enhancers and gene regulatory networks. Nat Methods 20, 1355–1367

85. Muerdter, F. et al. (2018) Resolving systematic errors in widely used enhancer activity assays in human cells. Nat Methods 15, 141–149

86. Shrikumar, A., et al. (2020) Technical Note on Transcription Factor Motif Discovery from Importance Scores (TF-MoDISco) version 0.5.6.5arXiv

87. Sundararajan, M. et al. (2017) Axiomatic Attribution for Deep Networks. arXiv

88. Armstrong, J. et al. (2020) Progressive Cactus is a multiple-genome aligner for the thousand-genome era. Nature 587, 246–251

89. Hickey, G. et al. (2013) HAL: a hierarchical format for storing and analyzing multiple genome alignments. *Bioinformatics* 29, 1341–1342

90. Cock, P.J.A. et al. (2009) Biopython: freely available Python tools for computational molecular biology and bioinformatics. Bioinformatics 25, 1422–1423

91. Litvin, U., et al. (2024) Viro3D: a comprehensive database of virus protein structure predictions. Cold Spring Harbor Laboratory

92. Kabsch, W. and Sander, C. (1983) Dictionary of protein secondary structure: pattern recognition of hydrogen-bonded and geometrical features. Biopolymers 22, 2577–2637

93. Joosten, R.P. et al. (2011) A series of PDB related databases for everyday needs. Nucleic Acids Res 39, D411–419

94. Jang, Y. and Bunz, F. (2022) AdenoBuilder: A platform for the modular assembly of recombinant adenoviruses. STAR Protoc 3, 101123

95. Jenner, R.G. et al. (2001) Kaposi’s sarcoma-associated herpesvirus latent and lytic gene expression as revealed by DNA arrays. J Virol 75, 891–902

