## Supplemental Figures for "Global cis-regulatory landscape of double-stranded DNA viruses"

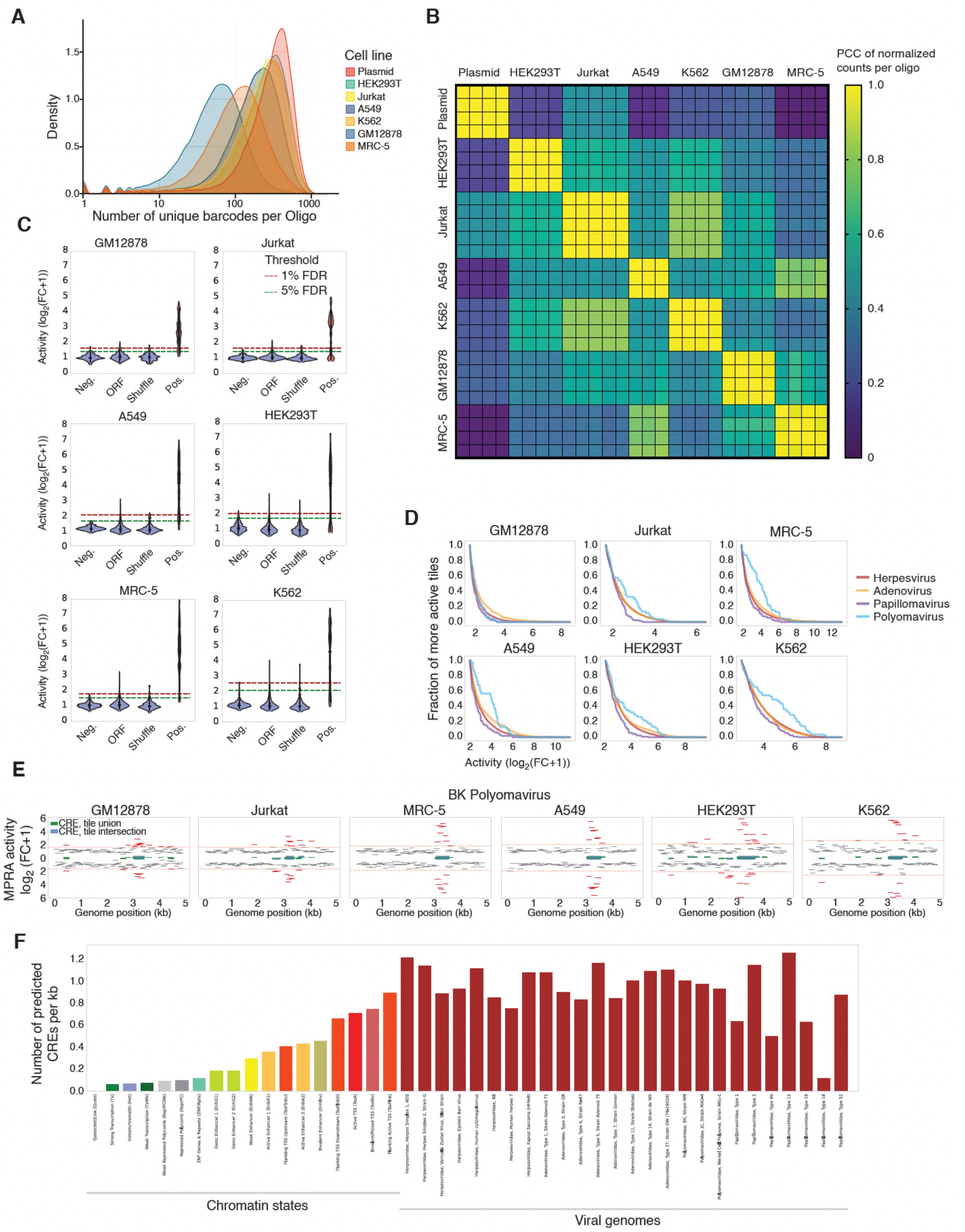
A
Cell line
Plasmid
HEK293T
Jurkat
A549
K562
GM12878
MRC-5
Density
Number of unique barcodes per Oligo
B
PCC of normalized counts per oligo
Plasmid
HEK293T
Jurkat
A549
K562
GM12878
MRC-5
C
GM12878
Jurkat
A549
HEK293T
MRC-5
K562
Threshold
1% FDR
5% FDR
Activity (log2(FC+1))
Neg.
ORF
Shuffle
Pos.
D
Fraction of more active tiles
Activity (log2(FC+1))
Herpesvirus
Adenovirus
Papillomavirus
Polyomavirus
E
BK Polyomavirus
CRE, tile union
CRE, tile intersection
MPRA activity log2 (FC+1)
Genome position (kb)
F
Number of predicted CREs per kb
Chromatin states
Viral genomes

**Figure S1: MPRA quality control of viral tiles, related to Figure 1**

(A) Cumulative distribution of MPRA oligos ranked by the number of unique barcodes (x-axis) detected across all replicates in the six cell lines and plasmid library. Red dashed line marks 10 barcodes.

(B) Correlation matrix of the summed barcode counts for each oligo across replicates and cell lines. Lower left matrix is a scatter plot of normalized oligo counts. A best fit linear regression is provided for each plot as a solid red line. Upper right matrices are Pearson correlations (r) for each pairwise comparison.

(C) MPRA activity distribution across cell lines for sequences known to be not active in MPRAs (Neg), and open reading frames (ORFs) and shuffled sequences, generally expected to be non-active, and positive controls with reported activity (Pos). Red and green dotted lines indicate 1% and 5% false discovery rate (FDR) thresholds.

(D) MPRA activity distribution in each cell line for active tiles of each viral family. The fraction of tiles more active than a particular activity level is indicated.

(E) CRE definition based on the union or intersection of consecutive active tiles per cell line. An example of BK Polyomavirus is shown.

(F) Number of predicted CREs per kb for each virus compared to those predicted within different chromatin states in the human genome.

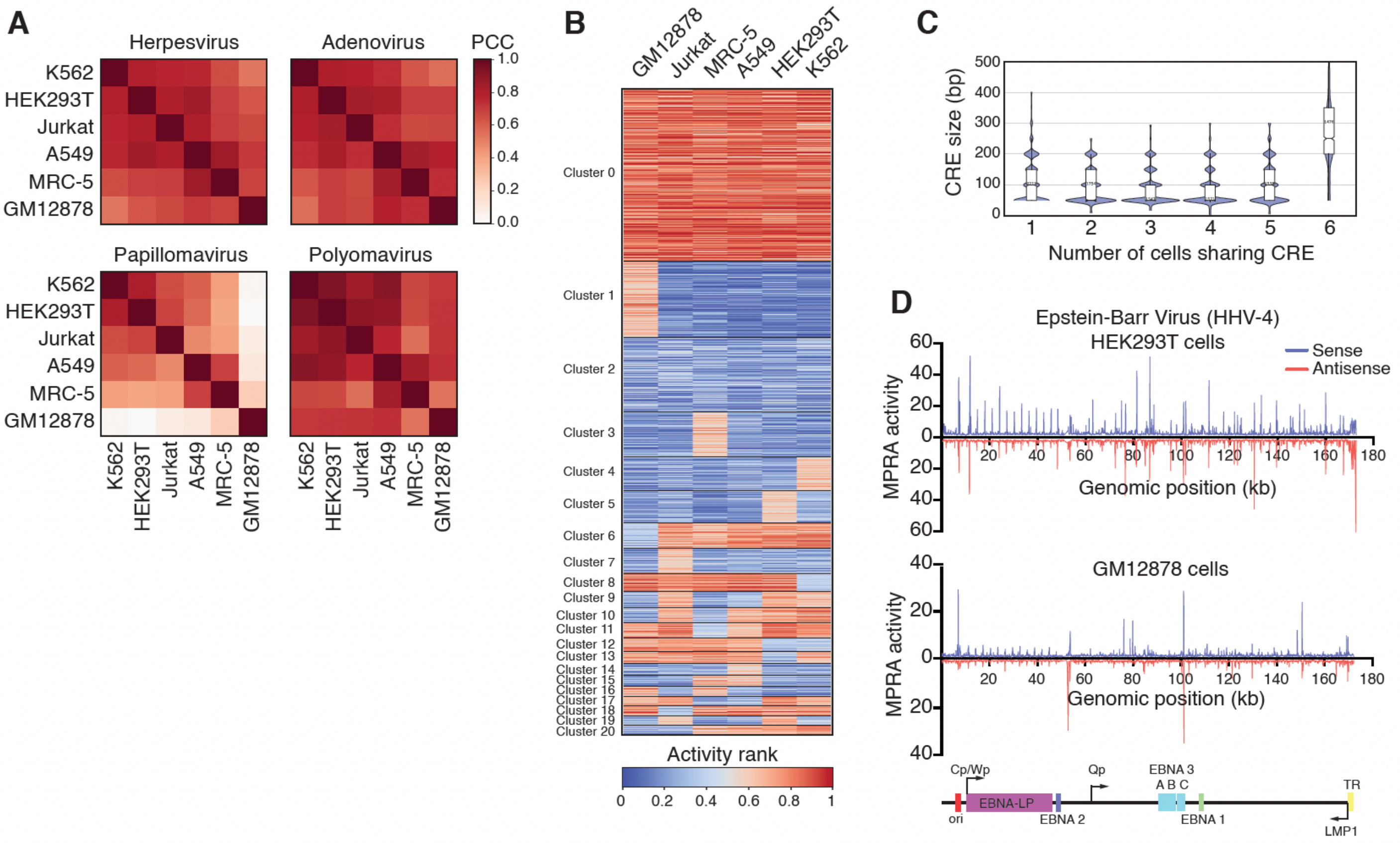


**Figure S2: Cell line specificity of viral tiles, related to Figure 1**

(A) Correlation (Pearson correlation coefficient) in activity for each viral family between cell lines across tiles active in at least one cell line.

(B) Activity ranks in each cell line for tiles active in at least one cell line. Tiles were clustered based on activity profiles.

(C) Distribution of CRE size, determined as the intersection of active regions across cell lines, based on the number of cell lines in which the CRE is active.

(D) MPRA activity map for HHV-4 in HEK293T and GM12878 cells. In blue and red is the activity when tiles were in positive and negative strand orientation, respectively.

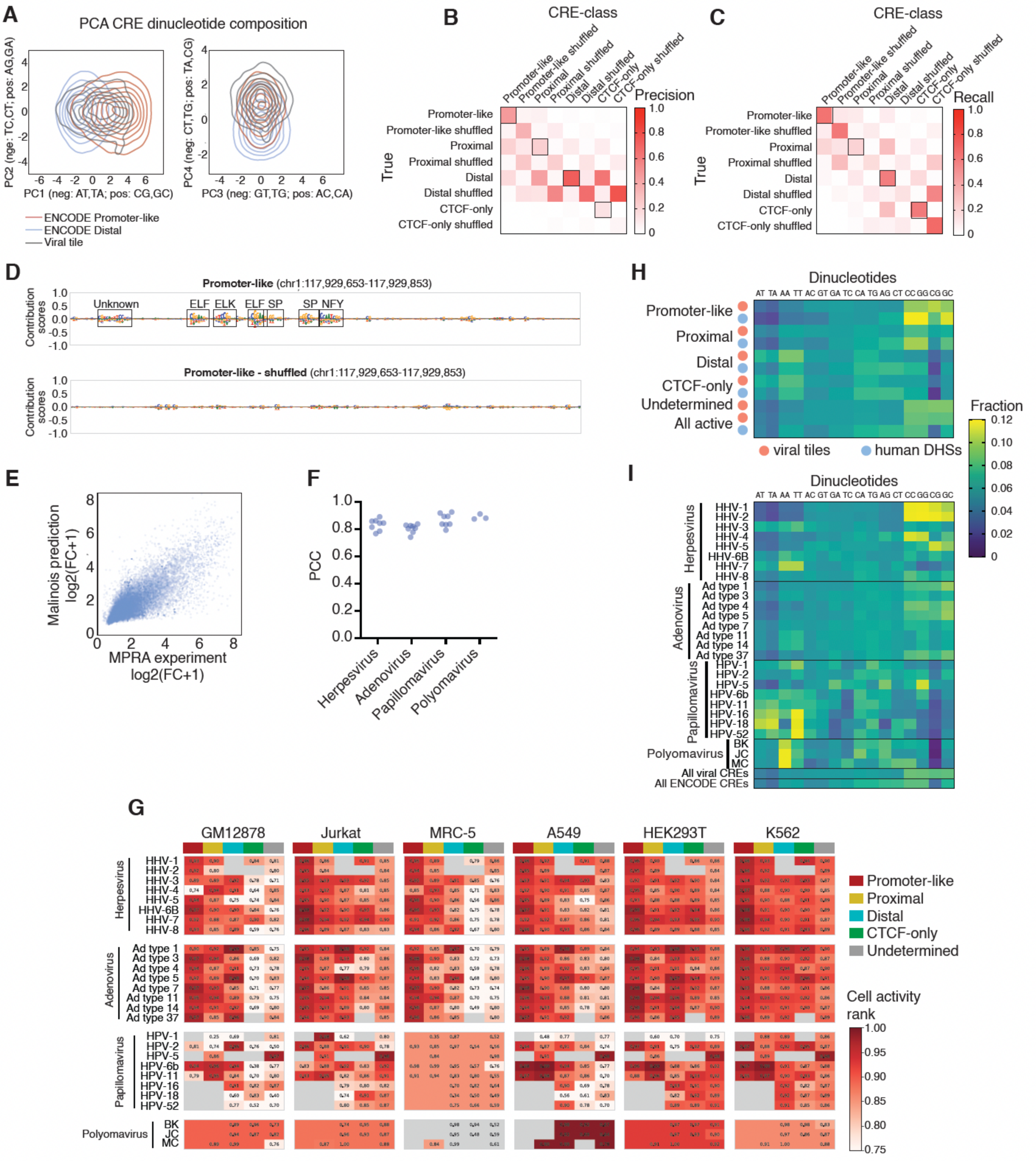


**Figure S3: Classification of viral CREs, related to Figure 2**

(A) PCA plots of tile dinucleotide composition for ENCODE promoter-like, ENCODE distal, and viral CREs. Density distribution is shown as contour maps.

(B-C) Precision (B) and Recall (C) o CRE-class to predict ENCODE CRE classes, as well as the corresponding shuffled sequences.

(D) Base contribution scores and motifs predicted to contribute to tile classifications for a human promoter-like located in Chr1:117,929,653-117,929,853, and for the corresponding shuffled sequence.

(E) Correlation between tile activity measured by MPRAs in K562 cells and activity predicted using Malinois trained on K562 cells.

(F) Pearson correlation coefficient (PCC) between measured and predicted activity for tiles from each virus.

(G) MPRA activity rank per cell type for tiles classified as promoter-like, proximal, distal, CTCF-only, and undetermined for each virus.

(H) Dinucleotide frequencies in active viral tiles or human ENCODE elements from different classes.

(I) Dinucleotide frequencies in active viral tiles from different viruses.

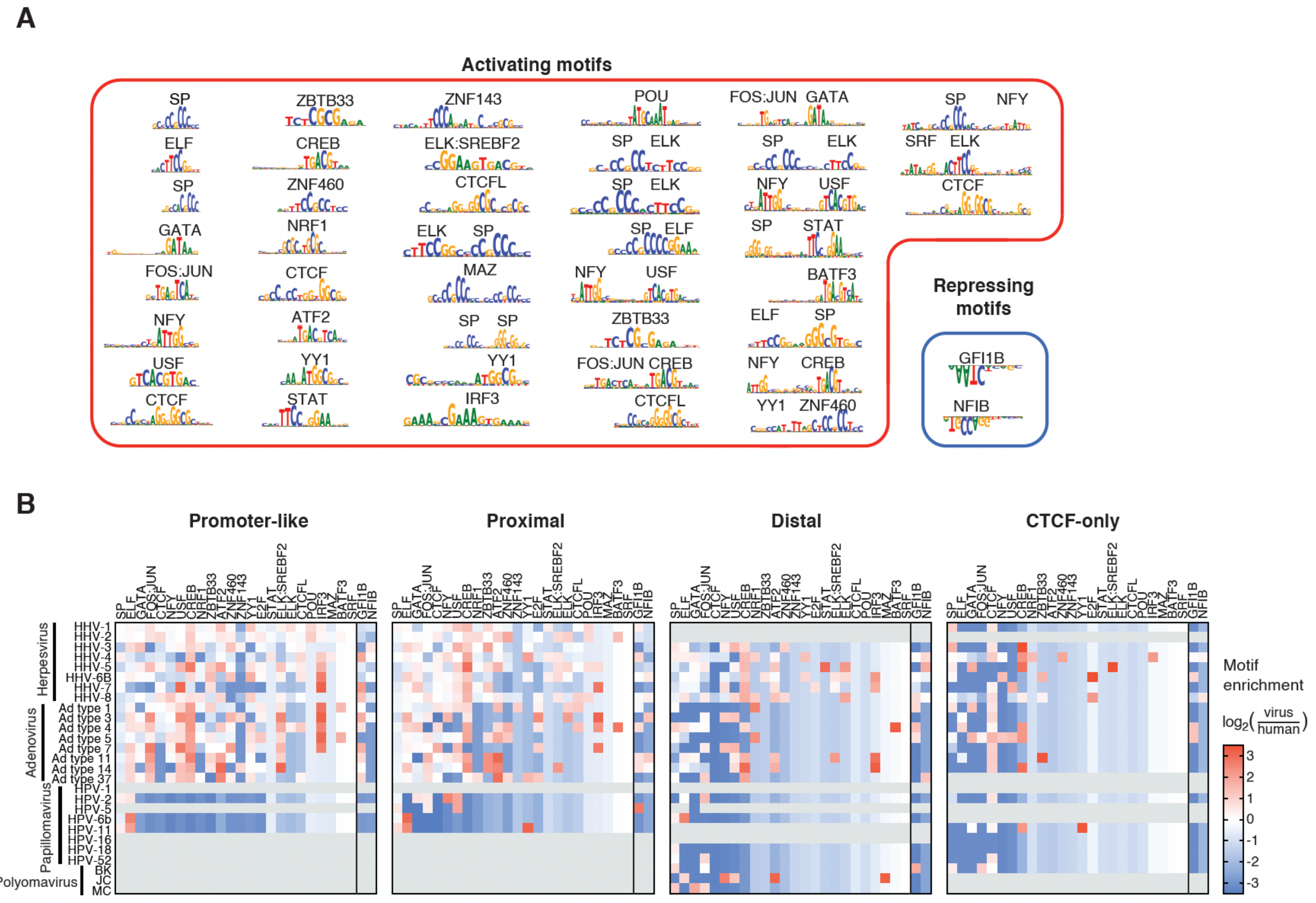


**Figure S4: TF modisco-lite motifs and frequency in active tiles, related to Figure 2**

(A) TF-modisco-lite motifs predicted to contribute to activity across viral tiles.

(B) Average number of motifs per active tile derived from TFmodisco-lite and collapsed into single TF motifs. Tiles are classified into promoter-like, proximal, distal, CTCF-only, and undetermined. TFs are ordered based on their frequency in human DHSs.

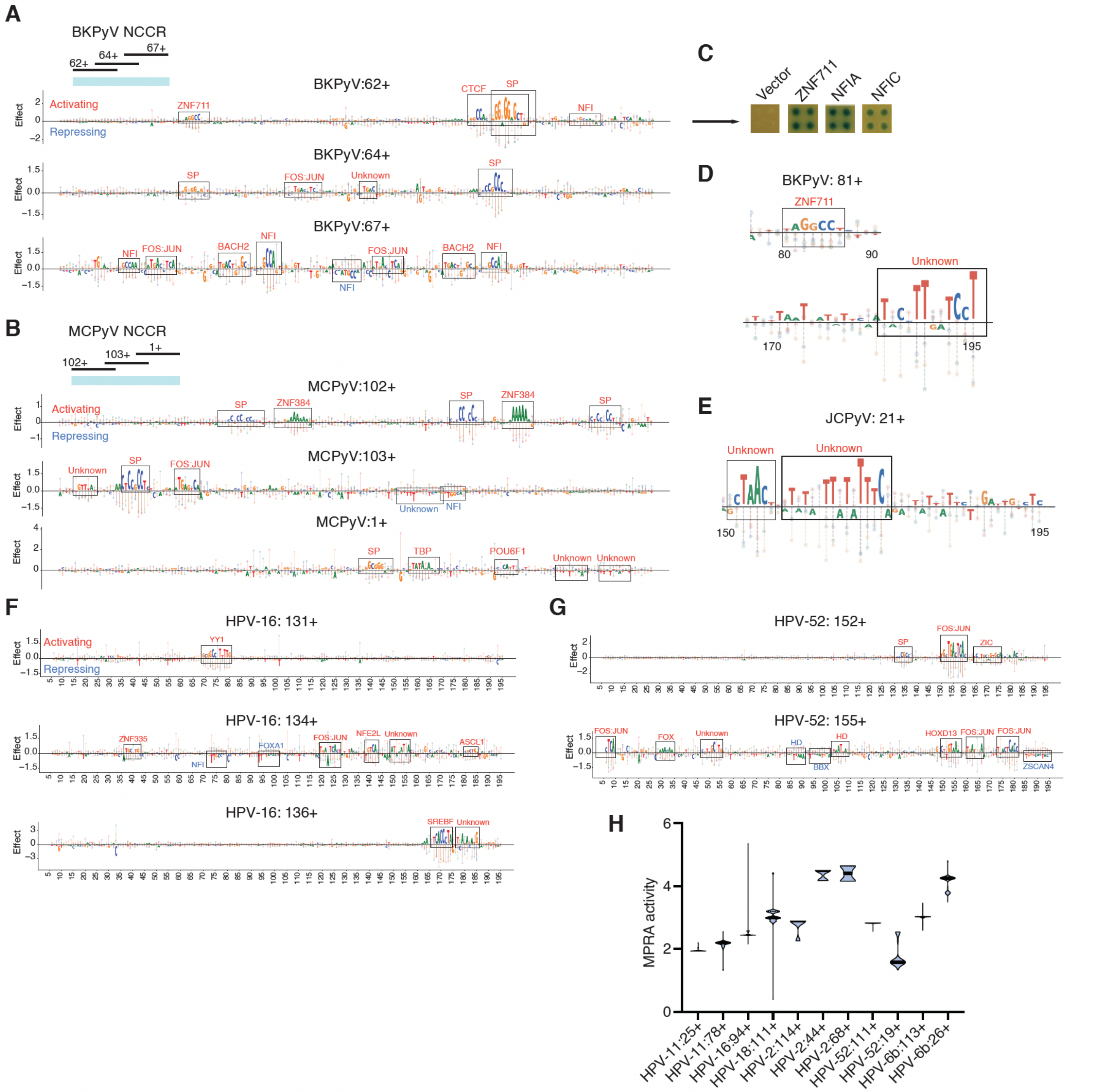


**Figure S5: Motif identification in Polyomavirus and Papillomavirus CREs, related to Figures 4 and 5**

(A-B) Base contributions scores derived from MPRA saturation mutagenesis results in HEK293T cells for tiles BK (A) and Merkel Cell (B) polyomavirus NCCRs. Activating and repressive motifs are indicated in red and blue, respectively.

(C) Enhanced yeast one hybrid assays showing interactions between the BK polyomavirus NCCR and 3 human TFs.

(D-E) Motifs examples from saturation mutagenesis experiments from tile 81+ from BK polyomavirus and tile 21+ from JC polyomavirus.

(F-G) Base contributions scores derived from MPRA saturation mutagenesis results in HEK293T cells for tiles HPV-16 (F) and HPV-52 (G) long control region. Activating and repressive motifs are indicated in red and blue, respectively.

(H) Violin plots of the tile activity measured by MPRAs in HEK293T cells for different isolates from different HPV types. The median and quartiles are indicated by bold solid and thin dashed lines, respectively.

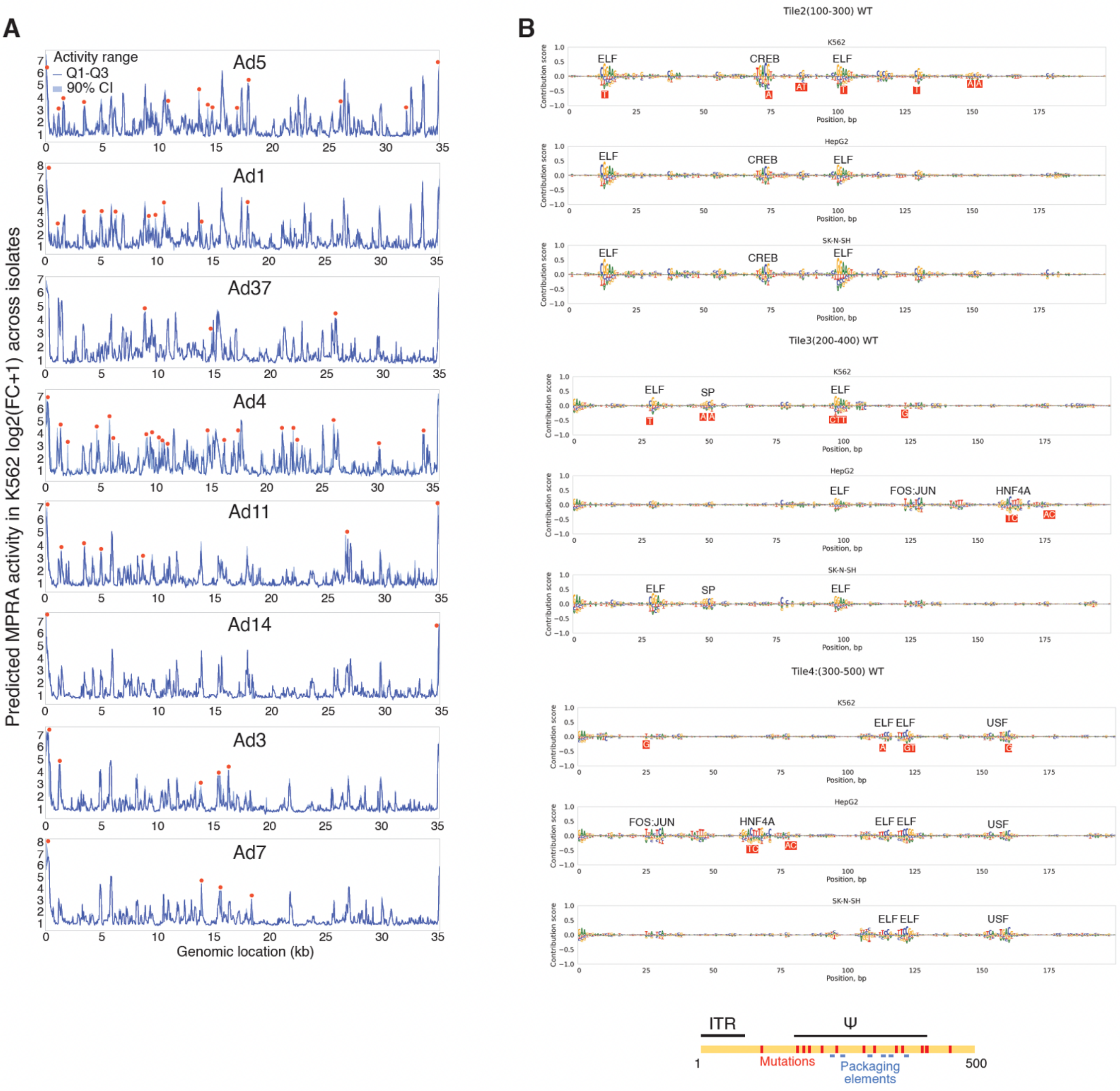


**Figure S6: Variation in Adenovirus CRE activity across isolates and editing of an adenovirus 5 vector**

(A) Activity levels, predicted using Malinois trained in K562 cells, across isolates of Adenovirus 5. Blue lines show the interquartile range, and light blue shade indicates the 90% confidence interval. Red dots show active regions for which the interquartile range is at least 2-fold.

(B) Base contributions predicted using Malinois trained in K562, HepG2, and SH-N-SH cells for tiles covering the Ψ region of the Adenovirus 5 vector. Red squares indicated the mutated bases.

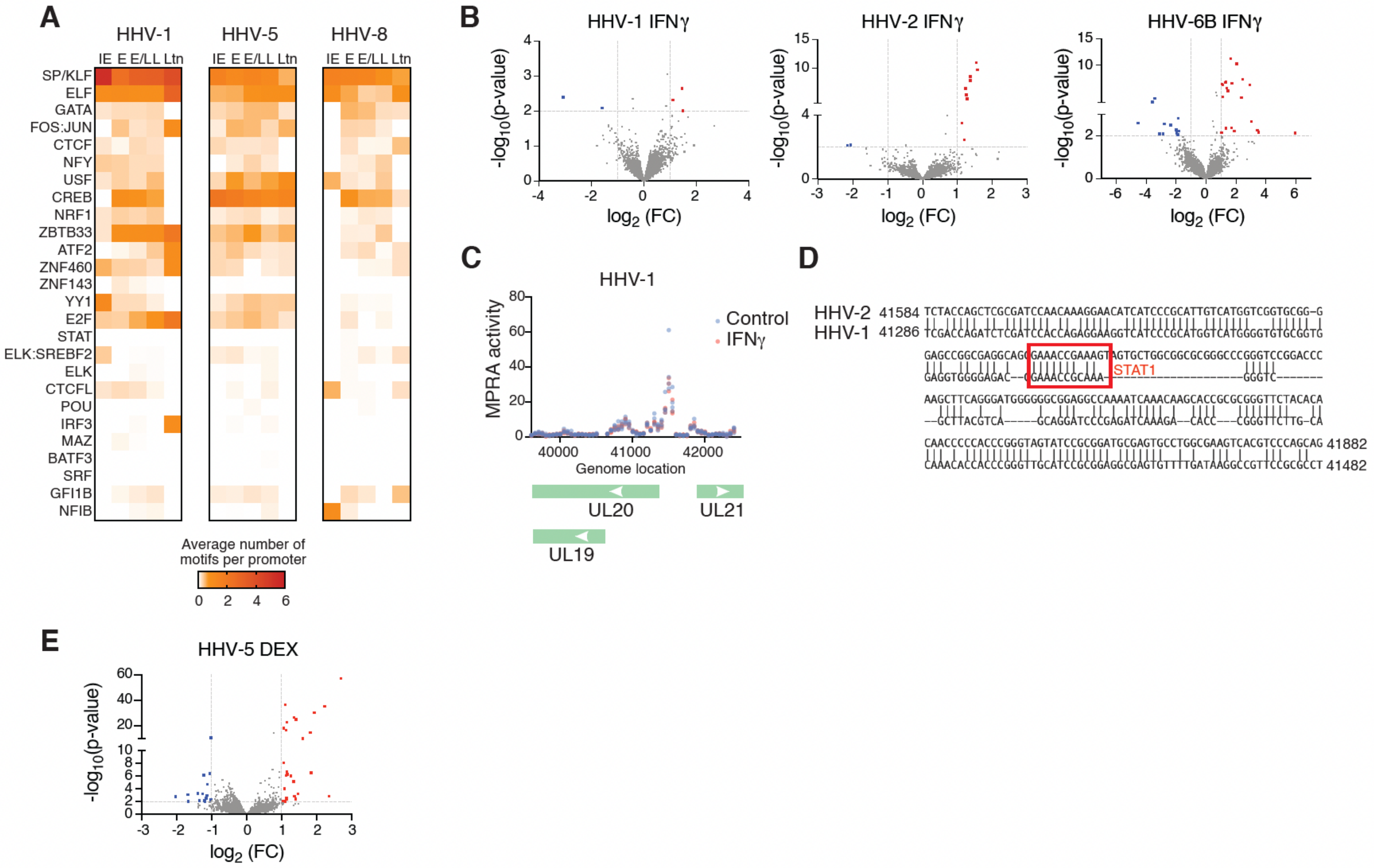


**Figure S7: Herpesvirus CREs motifs and activation, related to Figure 7**

(A) Heatmap of the average number of TF motifs in the promoters (±250 bp) of genes involved in different stages of viral replication: immediate early (IE), early (E), early/late (E/L), late (L), or latent (Ltn).

(B, E) Volcano plots showing the fold change in activity and significance for genomic sequence tiles in MPRA experiments in K562 cells stimulated with 100 ng/ml of IFN$\gamma$ versus control (B) or 100 nM dexamethasone versus control (E).

(C) Regions of HHV-1 activated by incubation of K562 cells with 100 ng/ml of IFN$\gamma$ for 6 hs.

(D) Alignment between HHV-1 and HHV-2 genomes highlighting a mutation in a STAT1 motif.
